# Nutrient availability influences *E. coli* biofilm properties and the structure of purified curli amyloid fibers

**DOI:** 10.1101/2023.09.07.556686

**Authors:** Macarena Siri, Mónica Vázquéz-Dávila, Cécile M. Bidan

## Abstract

Bacterial biofilms are highly adaptable and resilient to challenges. Nutrient availability can induce changes in biofilm growth, biomass, morphology, architecture and mechanical properties. Bacterial extracellular matrix plays a major role in achieving biofilm stability under different environmental conditions. Curli amyloid fibers are determining for the architecture and stiffness of *E. coli* biofilms, but how this major matrix component adapts to different environmental cues remains unclear.

Here, we investigated for the first time the effect of nutrient availability on both i) biofilm materials properties and ii) the structure and properties of curli amyloid fibers extracted from the biofilms. For this, we cultured *E. coli* W3110, which main matrix component is curli fibers. We quantified the size, mass and water content of the resulting biofilms and estimated their mechanical properties by microindentation. The curli amyloid fibers were then purified from the biofilms and their molecular structure and properties were studied by spectroscopic techniques. Our results show that the availability of nutrients in the substrate influences the yield of curli fibers, their structural composition and chemical stability, and suggest that these molecular features contribute to the stiffness of the biofilms. Biofilms grown on substrates with high nutrient concentration are softer, contain less curli fibers, and these fibers exhibit low β-sheet content and chemical stability.

Our multiscale study sheds new light on the relationship between the molecular structure of bacterial matrix and the macroscopic properties of biofilms. This knowledge will benefit the development of both anti-biofilm strategies and biofilm-based materials.

**Graphical abstract:** 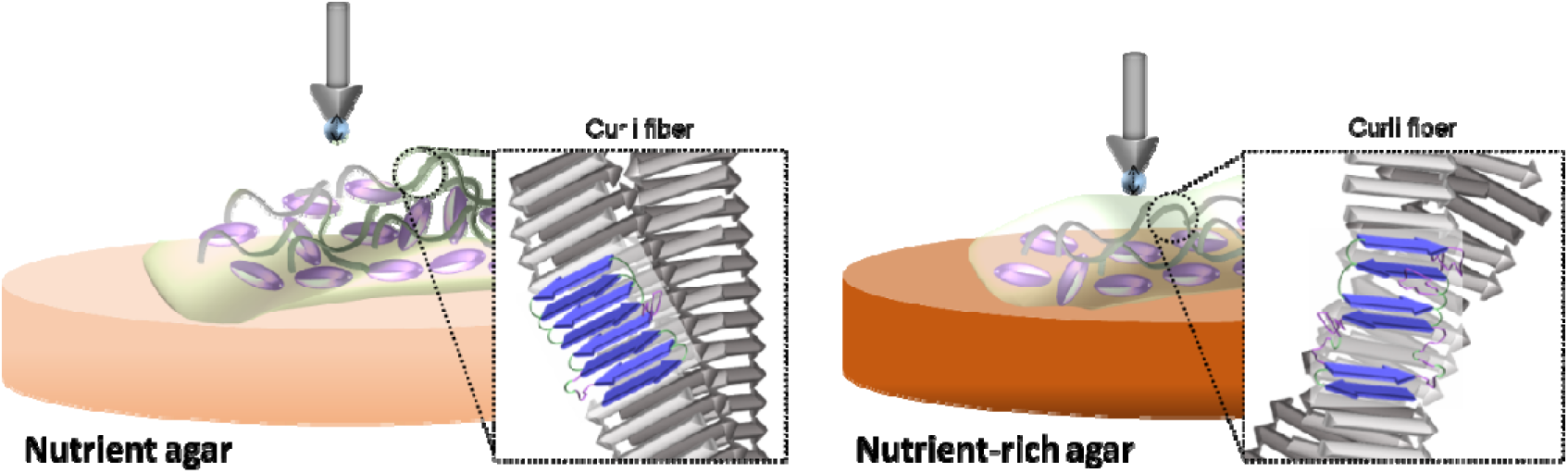

## Introduction

Bacteria co-exist in a community where they produce an extracellular matrix. Such living material is called a biofilm. Biofilm matrix typically contains DNA, exo-polysaccharides and proteins.^1^ Such ecosystems is a form of organization that, among other functions, protects the bacteria from environmental challenges. Biofilm properties thus depend on environmental physical cues such as temperature, pH and water availability.^2^

The availability of nutrients is also proposed to play a major role in the growth, phenotype, mechanical properties and matrix composition.^2–4^ For instance, nutrient limitations in the substrate might lead to spatial gradients of activities and molecules influencing the functioning of biofilms.^5^ Variation of nutrient availability and their effect on different biofilm aspects have been studied for different bacteria such as *Bacillus sp*.,^4,6^ *P. fluorescence*^7,8^ and *V. fischeri*^9^. The development and composition of biofilms can be influenced by nutrient availability through i) the variation of a specific nutrient,^9^ ii) the effect of metabolites in bacterial growth rate and mechanical properties^10^, iii) alteration of the environment by metabolites and iv) the addition of a substrate to promote the production of specific biopolymers by bacteria within the biofilm.^9–11^ Nevertheless, how these variations affect the molecular properties of the bacterial matrix and impact biofilm properties remains to be understood.

In amyloid-containing biofilms, like in *E. coli* and *P. aeruginosa* , amyloid fibers are known as the backbone of the biofilm as they provide structural integrity.^12^ In most cases, their molecular structure is influenced by the biofilm growth conditions.^7,13,14^ There is also evidence of a correlation between the biofilm macroscopic properties (physico-chemical and mechanical) and the molecular composition and structure of the material (interactions and entanglement between the matrix polymers and charged smaller molecules).^2,7,14^

Typically, amyloid fibers have a characteristic and stable quaternary conformation: cross-β structure with the β-strands arranged perpendicular to the fiber axis.^15,16^ Their structure and fibrillation mechanism have been studied throughout the years under *in vitro* conditions.^6,14,15,17^ Nonetheless, different studies describe not only an interaction between amyloids and components found in the biofilm matrix^13^, but also changes in the macroscopic features of the biofilm related to these protein assemblies.^3,7,12,14^ This kind of evidence allows new questions regarding how the natural environment in which amyloids are fibrillated contribute to determine their final structure.

This work focuses on the influence of nutrient availability on the structure of biofilm amyloid fibers and biofilm mechanics. We used the *E. coli* strain K12 W3110 that only produces curli fibers as the main component of its matrix, and cultured biofilms on agar substrates with different nutrient concentrations (i.e. yeast extract and tryptone). After 5 days, we quantified the size, mass and water content of the biofilms and estimated their mechanical properties by microindentation. Curli fibers were then purified from the biofilms, and fluorescence and ATR-FTIR spectroscopy were used to study the differences in their structure and functional properties like hydrophobicity and chemical stability. Overall, we provide knowledge to deepen the understanding of the relation between molecular features of the matrix components (e.g. fiber conformation) and macroscopic properties of the biofilm (e.g. stiffness).

## Results

Biofilm from *E. coli* K12 W3110 bacteria were grown on salt-free Lysogenic Broth (LB) agar substrates with different nutrient contents (0.75, 1.5, 3.0, 6.0 and 12.0 % w/v) (**Figure S1, Table S1** ). We analyzed the macroscopic features (i.e. size, mass, bacterial growth rate, total protein expression and biofilm stiffness) of the biofilms obtained in the different growth conditions. Curli fibers were then purified from the different biofilms and their structural and chemical characteristics were determined using spectroscopy techniques.

Of note, the agar composition containing 1.5 % nutrients is referred to as the standard biofilm growth condition because it has the same composition as the LB agar plates used to grow single bacterial colonies, without NaCl to promote biofilm formation.^18^

### Nutrient availability influences biofilm size, mass and rigidity

Variation in the nutrient availability of the agar substrate yielded biofilms different in size and morphology (**Figure 1a-b** and **Figure S2**). The area of biofilms increased from 220.2 ± 31.6 mm^2^ for substrates containing 0.75 % w/v nutrients, to 374.2 ± 15.8 mm^2^ for substrates containing 3.0 % w/v nutrients. The biofilm size decreased for biofilms grown on salt-free LB agar substrates containing higher nutrient content, reaching a size of 178.0 ± 34.7 mm^2^ on substrates containing 12.0 % w/v nutrients (**Figure 1b** and **Figure S2** ). This last condition also yielded biofilms different in shape (**Figure 1a** and **Figure S2**).

**Figure 1.**
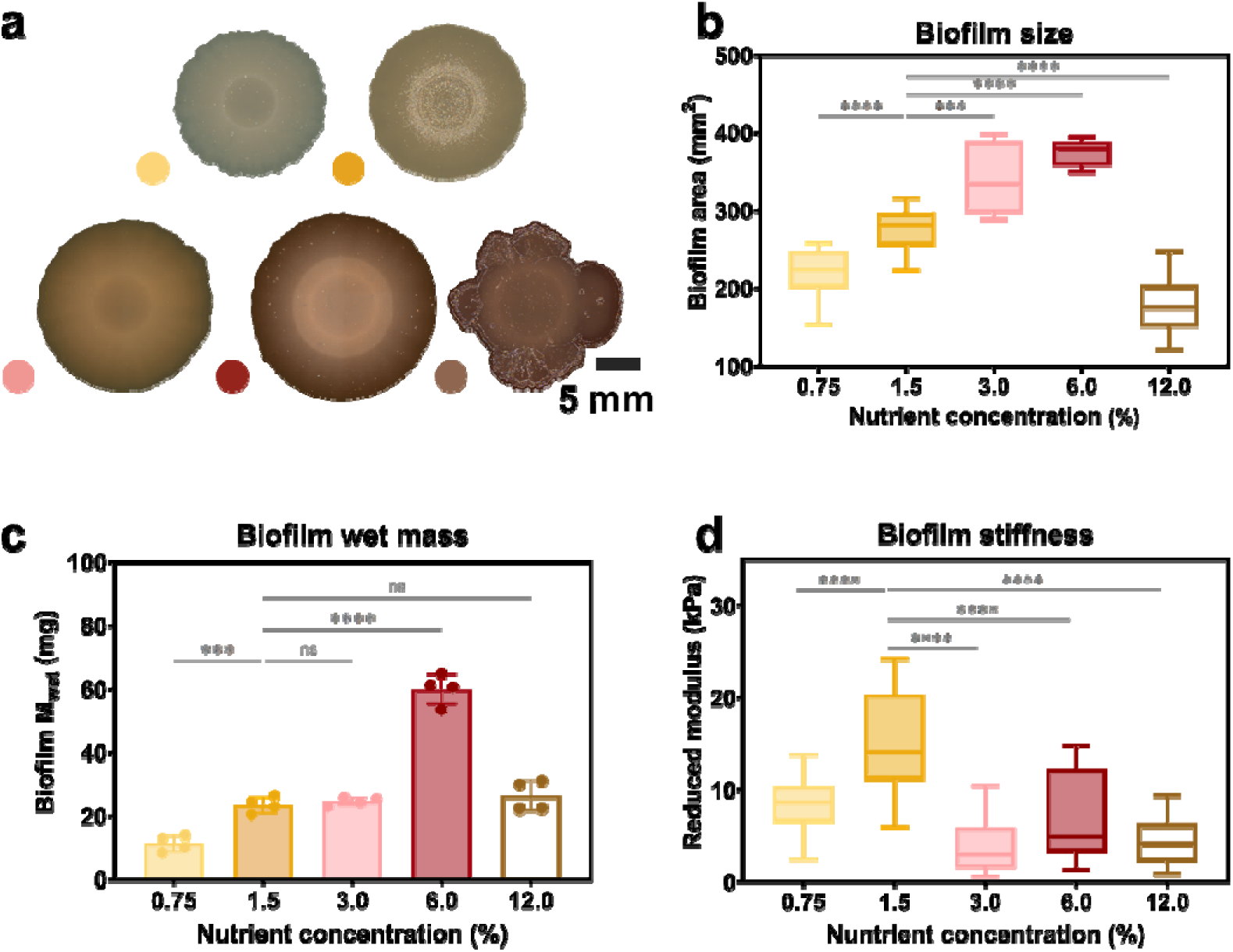
Macroscopic properties of *E. coli* W3110 biofilms agar substrates with various nutrient concentrations. (a) Representative phenotype of five-days old biofilms grown on substrates with different nutrient concentration. Scale bar= 5 mm. (b) Biofilm size based on their area. The statistical analysis was done with Man Whitney U test (p<0.0001, **** | p<0.001, *** | p<0.01, ** | p<0.05, * | ns = non-significant), where the 1.5 % w/v nutrient concentration condition was used as reference. Data acquired from 15 independent biofilms per condition. (c) Biofilm wet mass. Values are calculated for a single biofilm from four independent experiments. The statistical analysis was done with One-way ANOVA (p<0.0001, **** | p<0.001, *** | p<0.01, ** | p<0.05, * | ns = non-significant), where the 1.5 % w/v nutrient concentration condition was used as reference for the post-test multicomparisons. (d) Averaged reduced Young’s modulus obtained from microindentation experiments performed on biofilm surfaces. 10−23 individual measurements were done per condition. The statistical analysis was done with Man Whitney U test (p<0.00=1, **** | p<0.001, *** | p<0.01, ** | p<0.05, * | ns = non-significant), where the 1.5 % w/v nutrient concentration condition was used as reference for the post-test multicomparisons.

Along with their size, changes in the biofilm wet mass (M_wet_) were quantified by weighing (**Table 1**). We considered the *E. coli* W3110 biofilms wet mass (M_wet_) as the sum of the masses of i) the water, and ii) the bacteria and matrix, also considered as the dry mass (M_dry_). Within M_dry_, we discriminate between i) curli fibers (as the main matrix component) and ii) bacteria and other secondary matrix components. Biofilms grown on salt-free LB agar substrates containing 6.0 % w/v nutrients showed the highest wet mass, while biofilms grown at low nutrient availability (0.75 % w/v) presented the lowest wet mass (**Figure 1c** ). The rest of the conditions resulted in biofilms with similar wet mass. A similar trend was observed for the estimated dry mass in the biofilms (M_dry_) (**Figure S3** and **Table 1**).

**Table 1.** Composition of the *E. coli* W3110 biofilms grown on substrates of different nutrient concentration. The total mass corresponds to the sum of the water content and the dry mass. The bacteria mass was estimated by subtracting the curli mass given by the CsgA quantification after purification from the dry mass. The percentages of bacteria and CsgA are given with respect to the dry mass. N= 4 biofilms per experiment.

| Substrate nutrient concentration (% w/v) | Biofilm size (mm) | Units | Total mass (M<sub>wet</sub>) | Water | Dry mass | CsgA<sup>a</sup> | Bacteria<sup>a</sup> |
| --- | --- | --- | --- | --- | --- | --- | --- |
| 0.75 | 16.65 ± 1.49 | mg | 11.38 ± 2.60 | 8.35 ± 1.75 | 3.04 ± 0.86 | 0.01 ± 0.00 | 3.03 ± 0.86 |
|  |  | % | 100 | 73 | 27 | 0.35 | 99.65 |
| 1.50 | 18.19 ± 1.92 | mg | 23.56 ± 2.53 | 17.57 ± 2.00 | 6.01 ± 0.57 | 0.02 ± 0.01 | 5.99 ± 0.57 |
|  |  | % | 100 | 75 | 26 | 0.36 | 99.64 |
| 3.00 | 19.91 ± 2.06 | mg | 24.64 ± 1.28 | 18.38 ± 0.96 | 6.29 ± 0.32 | 0.03 ± 0.01 | 6.26 ± 0.32 |
|  |  | % | 100 | 75 | 26 | 0.50 | 99.50 |
| 6.00 | 19.76 ± 1.59 | mg | 60.04 ± 4.79 | 40.24 ± 4.26 | 19.86 ± 0.96 | 0.05 ± 0.01 | 19.80 ± 0.97 |
|  |  | % | 100 | 67 | 33 | 0.28 | 99.72 |
| 12.00 | 12.85 ± 0.91 | mg | 26.53 ± 4.74 | 17.63 ± 3.05 | 8.90 ± 1.69 | 0.01 ± 0.00 | 8.89 ± 1.69 |
|  |  | % | 100 | 66 | 34 | 0.10 | 99.90 |

**Table 2.** FWHH input values in cm^−1^ and their physically plausible ranges expected for each type of secondary structure^16,20,21^.

| Secondary structure component | | FWHH input<br>($\text{cm}^{-1}$) | Lower limit<br>($\text{cm}^{-1}$) | Upper limit<br>($\text{cm}^{-1}$) |
| --- | --- | --- | --- | --- |
| β-sheet | High wavenumber component | 9 | 8 | 11 |
|  | Low wavenumber component | 17 | 14 | 19 |
| α-helix/ random | | 20 | 5 | 30 |
| Turns |  | 20 | 5 | 30 |

We carried out microindentation experiments to study whether these changes in size and wet mass had an effect on the stiffness of the biofilms (**Figure 1d** and **Figure S4** ). The biofilm stiffness was estimated by indenting up to 10 μm in the center of the biofilm (10 % of its thickness approximately) to avoid influence of the substrate during the measurement (**Figure S4, inset**). Biofilms grown on substrates containing 1.5 % w/v nutrients presented the highest stiffness value, 15.28 ± 5.07 kPa (**Figure 1d**). Biofilms grown on substrates containing 3.0 and 12.0 % w/v nutrients showed the lowest values with 3.71 ± 2.78 and 4.26 ± 2.42 kPa, respectively. Results suggest that the stiffness from the biofilm is independent of their size or their wet mass.

### Nutrient availability influences biofilm composition and ability to uptake water

Biofilms were further characterized to estimate their composition, including their water content and their water uptake ability (**Figure 2** and **Table 1**). The water content was assessed by weighing the biofilms before and after dehydration. Biofilms grown at low nutrient availability (0.75 – 3.0 % w/v) presented the highest water content c.a. 74 % w/w, while biofilms grown at high nutrient availability presented the lowest water content, approximately 66 % w/w (**Figure 2a** ). The water uptake ability of the biofilms was tested upon overnight rehydration (**Figure 2b**). Except for biofilms grown on salt-free LB agar containing 3.0 and 6.0 % w/v nutrients, all biofilms presented a similar rehydration behavior. Biofilms grown at low nutrient availability (and at 12.0 % w/v nutrient concentration) absorbed 55 – 70 % more of water than their original weight. Biofilms grown on salt-free LB agar containing 3.0 and 6.0 % w/v nutrients absorbed 30 % of water more than their original weight. Analysis of the water uptake per gram of dry biofilm after rehydration showed that, the lower the nutrient availability, the higher the water uptake (**Figure 2c**).

**Figure 2.**
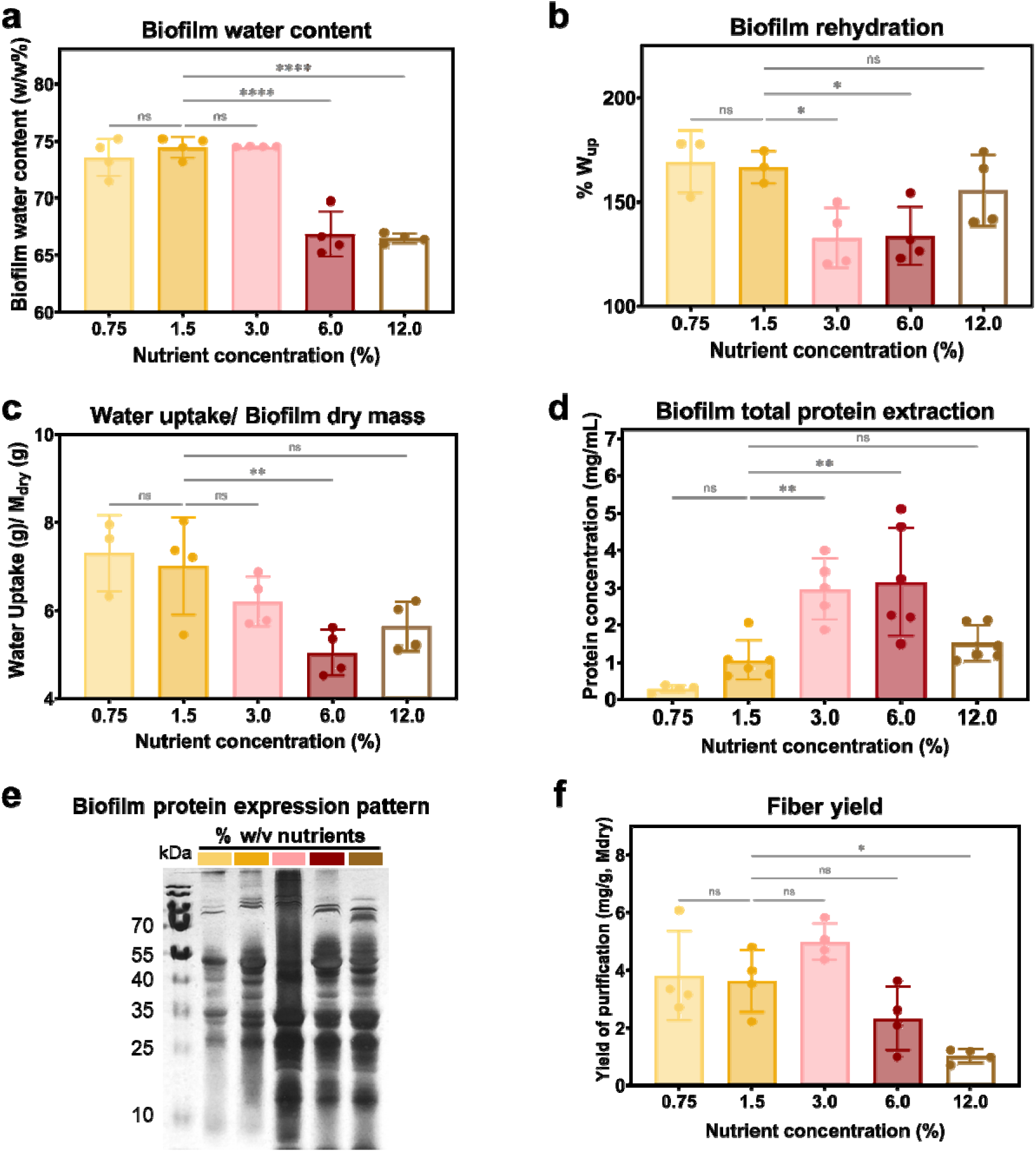
General characteristics of the composition of *E. coli* W3110 biofilms grown on substrates with different nutrient contents. (a) Biofilm water content. (b) Biofilm water uptake upon overnight rehydration of dry mass. (c) Biofilm water uptake per gram of dry mass. (d) Biofilm total protein concentration (see experimental section for details). (e) SDS-PAGE electrophoresis gel with showing the protein pattern of the samples studied in (d). (f) Purification yield of the curli fibers extraction process in milligram of CsgA per gram of dry mass. All data presented here come from N=4 independent biofilm cultures for each condition tested. The statistical analysis was done with One-way ANOVA (p<0.0001, **** | p<0.001, *** | p<0.01, ** | p<0.05, * | ns = non-significant), where the 1.5 % w/v nutrient concentration condition was used as reference for the post-test multicomparisons.

To deepen our understanding on how the biofilm matrix changes with biofilm growth conditions, we focused on the biofilm composition and first estimated the total amount of proteins using a Bradford assay (see Experimental section for more details). Biofilms grown on salt-free LB agar containing 3.0 and 6.0 % w/v nutrients had the highest total protein concentration (**Figure 2d**). The trend observed for the biofilm protein concentration is similar to the one observed for the size and wet mass of the biofilms, as well as for the dry mass of the biofilms (**Figure S3** ). The SDS-PAGE electrophoresis gel run with aliquots of each total protein extraction showed that the pattern of proteins expressed by the bacteria is similar between conditions (**Figure 2e**).

To clarify whether these differences in protein production are due to the growth and the metabolism of the bacteria, we monitored the bacteria growth for 8 hours in liquid salt-free LB media and calculated their doubling time (T_d_) (**Figure S5-S6** ). While the growth curves and T_d_ was similar for all conditions, the final optical density of the bacteria increased with the nutrient availability in the media (**Figure S6** ). The bacteria metabolic activity was also studied in each phase of their growth curve (**Figure S6** ). *E. coli* bacteria in liquid salt-free LB media containing 12 % w/v nutrients were more metabolically active from the beginning of the exponential phase compared to the other nutrient concentrations tested.

Among the proteins present in the biofilm, curli amyloid fibers are the main components in the biofilm matrix. We purified them and estimated a yield by quantifying the monomeric unit (CsgA) and normalizing per gram of dry biofilm (**Figure 2f** ). Interestingly, biofilms grown on salt-free LB agar substrates containing 12.0 % w/v yield two times less purified curli fibers compared to the other conditions. No significant differences were observed for the other yields in fiber production. We observe that the curli fibers yield does not follow the trend of the biofilm total protein concentration, nor the trend of the bacterial metabolic activity. However, there is a tendency for increased biofilm water uptake at higher fiber yields. (**Figure S7**).

### Nutrient availability influences the packing and structure of curli amyloid fibers in the biofilms

The structure of curli fibers after the purification process was examined by transmission electron microscopy (TEM), by Thioflavin T fluorescence emission (ThioT) and by Fourier transform infrared (ATR-FTIR) spectroscopy (**Figure 3a-c**). TEM confirmed the presence of curli fibers in the purified product (**Figure S7** ). Following basic amyloid fiber characterization, we stained the fibers with ThioT. This dye undergoes a fluorescence enhancement when immobilized in the β-sheets of the amyloid fibers due to binding.^19^ Despite every sample having ThioT fluorescence emission, there were differences in the intensity values (**Figure 3a**). When normalizing the intensity value against the free ThioT signal, fibers produced on substrates containing 1.5 and 3.0 % w/v of nutrient showed an increase of c.a. 15 times of the ThioT intensity (**Figure 3b**). Fibers produced in a context of higher or lower nutrient availability showed only a five-fold increase in the ThioT intensity. The differences observed here suggest differences in the packing or in the β-sheet content of the fibers.^14^

**Figure 3.**
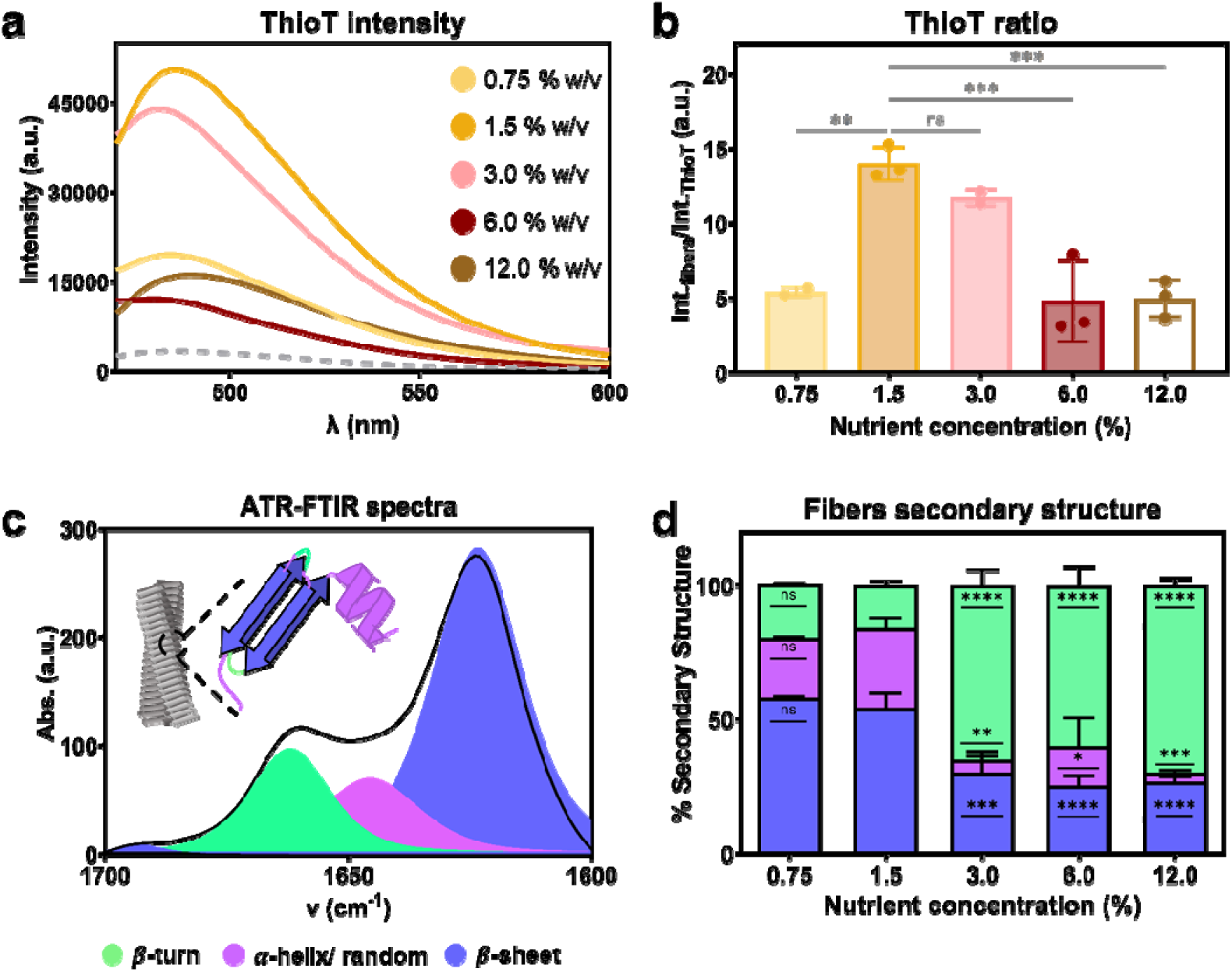
Characterization of the structure of purified curli fibers. (a) ThioT fluorescence emission spectra of the fibers at equivalent mass concentrations. (b) Values describing the increase in intensity of ThioT when bound to the purified fibers. Quantification of the increase was estimated by division of the area under the curve of each spectra of the probe with each fiber by the area under the curve of the emission spectra of the probe alone. The statistical analysis was done with One-way ANOVA (p<0.001, *** | p<0.01, ** | p<0.05, * | ns = non-significant), where the 1.5 % w/v nutrient concentration condition was used as reference for the post-test multicomparisons. (c) Representative Amide I’ region of an ATR-FTIR spectrum for curli fibers. Each peak assigned to the different secondary component is highlighted in their respective color. (d) Distribution of the three types of secondary structure in the curli fibers obtained in the different growth conditions. The data was obtained from the Amide I’ region of each spectra. The statistical analysis was done with One-way ANOVA (p<0.0001, **** | p<0.001, *** | p<0.01, ** | p<0.05, * | ns = non-significant), where the 1.5 % w/v nutrient concentration condition was used as reference for the post-test multicomparisons.

ATR-FTIR spectroscopy was used to further study the fiber structure (**Figure 3c-d**). We focused on the amide I1. band (from 1700 to 1600 cm^−1^), i.e. the fingerprint region for protein analysis (**Figure 3c** ).^20–22^ All samples showed spectra expected for curli amyloid fibers (**Figure 3c** and **Figure S8**): a band around 1620 cm^-1^ assigned to P-sheet structures, and second band around 1660 cm^-1^ usually assigned to P-turns structures.^14,15^ The intensity and width of the band at 1660 cm^-1^ increased with the nutrient availability, indicating differences in the secondary structure composition of the fibers (**Figure S8**). We used a band fitting method to decompose the spectra into the secondary structure components (**Table 1** ).^14,20^ Purified curli fibers obtained from biofilm grown at low nutrient availability (0.75 and 1.5% w/v) had the highest content of β-sheet and the lowest content of β-turn structures (**Figure 3d** and **Table S3** ). In contrast, purified curli fibers obtained from biofilm grown at higher nutrient availability (3.0 – 12.0 % w/v) had the lowest content of β-sheet and the highest content of β-turn structures (**Figure 3d** and **Table S3** ). Differences in the β-sheet structure of the fibers was also observed by CD spectroscopy (**Figure S9** ). These results do not follow the trend presented in the ThioT experiment, suggesting that the latter does not mainly report differences in β-sheet content. Thus, in this case, differences in ThioT intensity mainly report differences in fiber packing.

### Fiber packing and structure determine curli physico-chemical features

Since the structure of a molecule is often related to its properties and functions, we explored if the differences observed on the fiber conformation are reflected in their physico-chemical features. The global hydrophobic character of the purified curli fibers was studied by binding to Nile Red (NR) (**Figure 4a** and **S10a** ). NR is a solvatochromic dye reporting the hydrophobic character of the molecule it binds to. As the emission spectrum of NR in buffer is different from the emission spectrum of NR in ethanol, we used these spectra as references to assess the hydrophobic character of the amyloid fibers obtained in the different conditions (**Figure 4a**). All purified fibers present a strong hydrophobicity. However, as the nutrient availability in the substrate increased, the hydrophobic character of the fibers also increased (**Figure S10a**). Curli fibers are made of CsgA monomers, each of them containing a single tryptophan (Trp). Trp is an amino acid behaving as a solvatochromic fluorophore (**Figure S10b**),^14^ which position of maximum fluorescence emission gives information on the hydrophobicity of its environment (**Figure 4b, inset** ).^23^ Curli fibers grown at a low nutrient concentration (0.75 % and 1.5 % w/v) presented a Trp population in a less hydrophobic environment than the other fibers.

**Figure 4.**
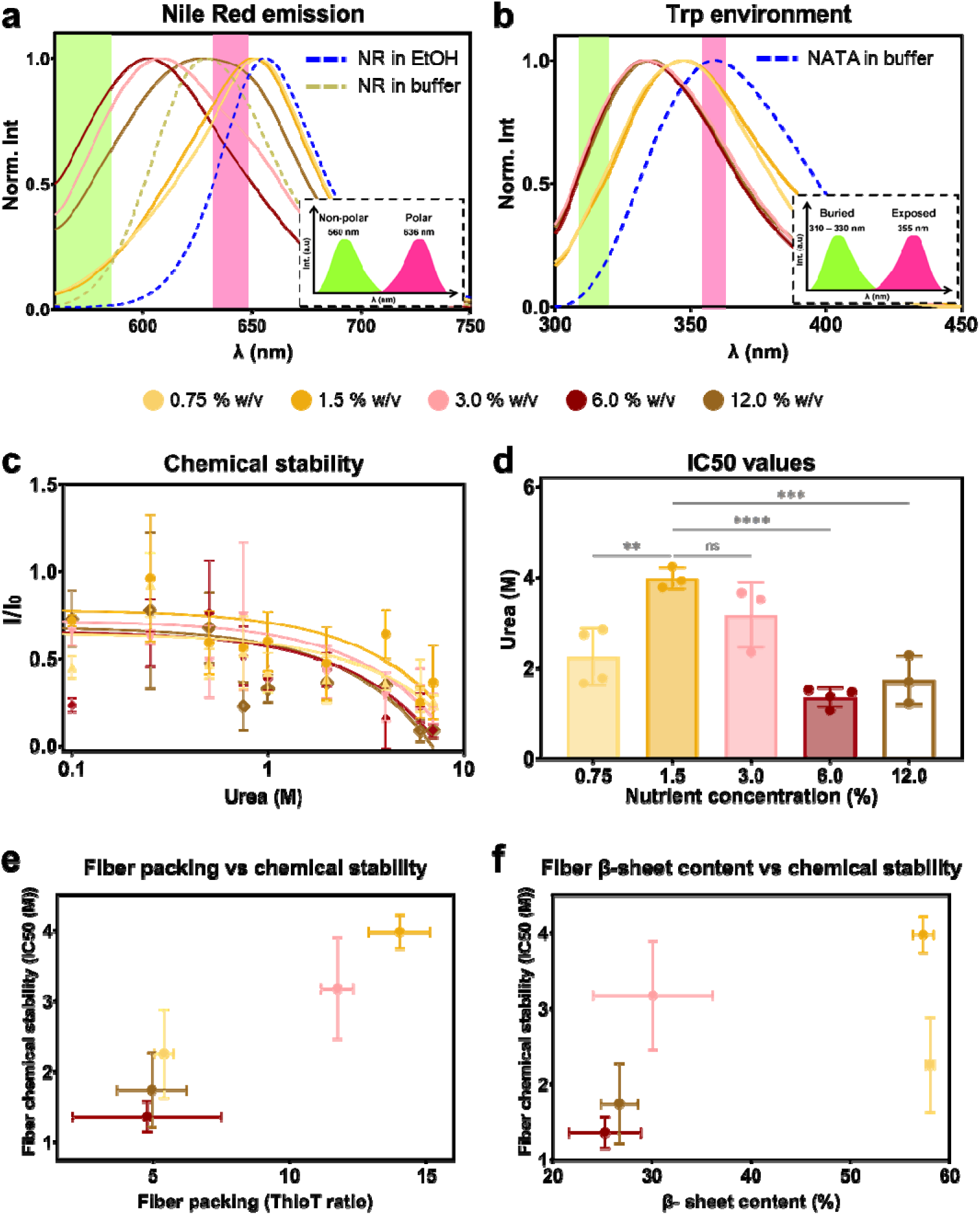
Structure-function features of the purified curli fibers. (a) Fluorescence spectra of fibers stained with Nile Red (NR). The spectra of NR in ethanol and in buffer are represented as references for the emission of NR in a hydrophobic and hydrophilic environment, respectively. A sketch of the position of peaks for the NR in non-polar and polar environments is represented in the inset^24^. The shadowed areas in the main plot indicate these exposure extremes. N=3-4 independent biofilm cultures for each condition tested. (b) Intrinsic fluorescence of the fibers through Trp emission. The spectrum of soluble Trp (NATA) in buffer is represented as reference for the maximum exposure possible of the Trp to the surface (λ_exc_ = 280 nm). A sketch of the position of peaks for buried and exposed Trp is represented in the inset^25^. The shadowed areas in the main plot indicate these exposure extremes. N=3-4 independent biofilm cultures for each condition tested. (c) Chemical stability of the purified fibers upon denaturation with increasing urea concentrations (0.1 – 8 M). The presence of the fibers was observed by ThioT fluorescence emission intensity. I_0_ corresponds to the ThioT emission when bound to fibers without urea in the solution. (d) IC50 values for each curve in panel (c) IC50 corresponds to the urea concentration at which the ThioT intensity is 50 % of the initial one. (e) Relationship between fiber packing (ThioT ratio) and their chemical stability. (f) Relationship between fiber β-sheet content and their chemical stability. N=6 independent fiber samples. The statistical analysis was done with One-way ANOVA (p<0.0001, **** | p<0.001, *** | p<0.01, ** | p<0.05, * | ns = non-significant), where the 1.5 % w/v nutrient concentration condition was used as reference for the post-test multicomparisons.

Finally, we studied the implication of the different structures of purified curli fibers by testing their chemical stability (**Figure 4c-d** ). Fiber denaturation into CsgA monomers was monitored by the fluorescence intensity emitted by ThioT.^14^ Curli fibers obtained from biofilms grown on salt-free LB agar containing 1.5 % w/v nutrients had the highest stability, while fibers grown at lower nutrient availability (0.75 % w/v) or higher than 3.0 % w/v presented low chemical stability (**Figure 4c**). Interestingly, fibers with a high packing degree and a high β-sheet content, presented high chemical stability (**Figure 4e-f**). The higher the packing in the fiber, the higher the chemical stability. While biofilm stiffness may be influenced by different fiber properties, no further particular trend emerged (**Figure S11**).

## Discussion

In the present work, we studied how nutrient availability influences *E. coli* biofilms from the macroscopic to the molecular scale (**Figure 5** ). The results show that biofilm size, mass and rigidity change as the amount of surrounding nutrients varies (**Figure 1**). Such effects of yeast extract and tryptone variations on macroscopic biofilm properties (e.g. morphology, thickness and density) have been reported for other bacteria strains.^4,9,26^ In the case of *E. coli* K12 W3110, we observed that the relation is not monotonic and rather shows a maximum of these properties at intermediate nutrient availability, namely 6 % w/v for the size and mass and 1.5 % w/v for the stiffness (**Figure 1**). The existence of such optima may result from simultaneous and opposite effects of substrates properties that cannot be decoupled from the nutrient content. For example, both the presence of nutrients and water are expected to promote biofilm growth respectively *via* the stimulation of bacterial activity, the lubrication of the surface and biofilm swelling,^26,28^ yet salt-free LB agar containing higher amounts of nutrients also happen to contain less free water (**Figure S1 a-c** ). Nutrient availability has also proven to be as important at the initial steps of the biofilm formation e.g. in the layout of the bacteria,^29^ as at the later stages in which water channels form for nutrient distribution,^11^ and gradients are established.^30^ The implication of nutrients at different hierarchical levels of the biofilm architecture can thus explain the effect of their concentration on a broad range of biofilm properties.^31^

**Figure 5.**
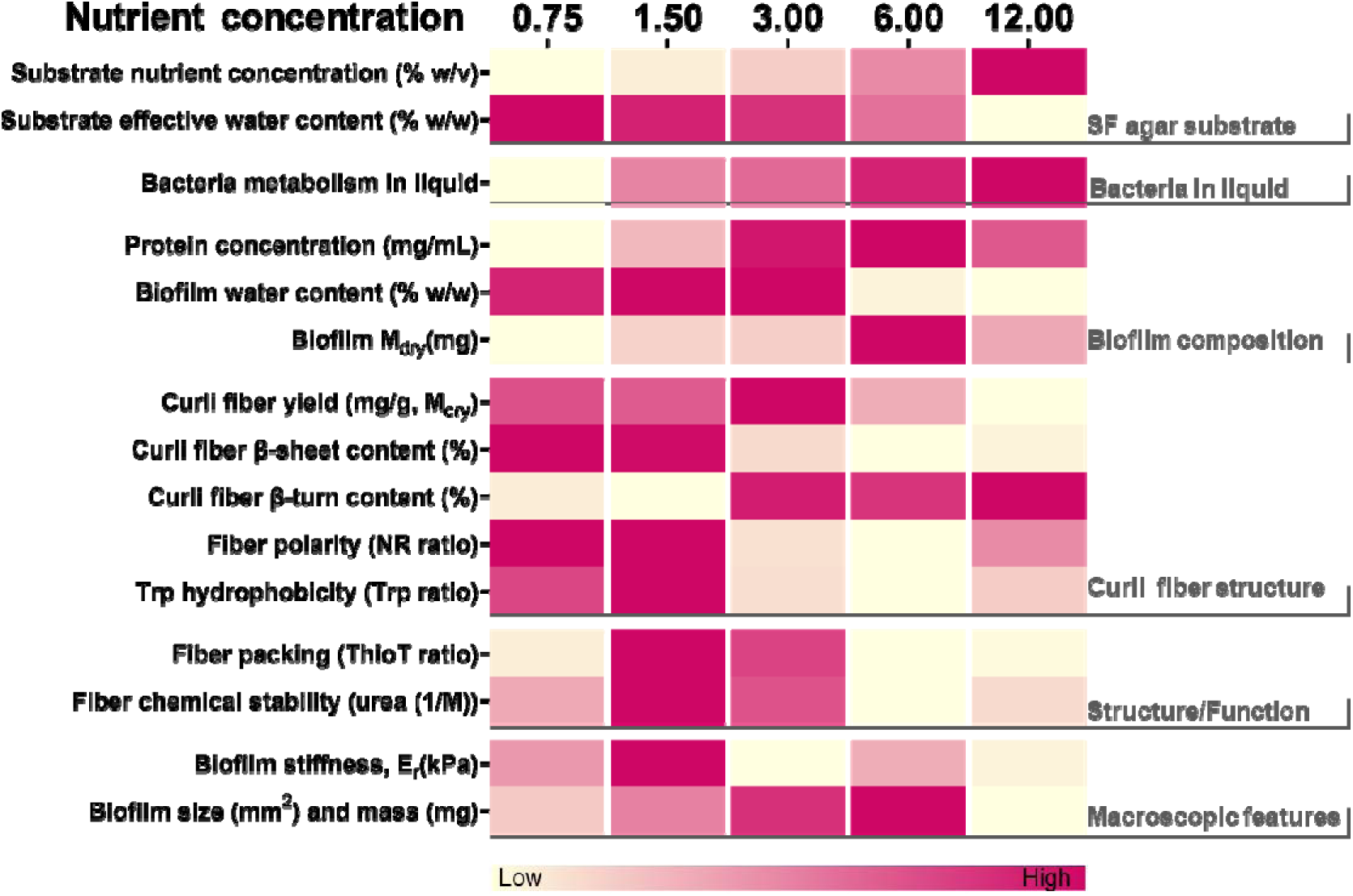
Summary of the relationship between nutrient variation in the salt-free LB agar and the macro- and microscopic features of the biofilms grown under each condition. The figure serves as a visual global analysis of the data acquired and guide to the following discussion integrating the results obtained in this work. We indicate color shade the position of the parameter value in the column on the left of one of the conditions compared to the other conditions studied (yellow for the lowest value, to fuscia for the highest value).

As we decomposed the biofilm mass obtained into the contributions of water, bacteria and curli fibers, we observed that biofilm water content and curli fiber yield both tend to be higher when nutrients are less abundant (**Figure 22a** and **2f** ). Although not the sole factor, amyloid fibers are known to contribute to keeping a hydrated microenvironment within biofilms.^32^ This can be explained by the potential of amyloid fibers to take up and bind water (**Figure 2b and c**).^33^ Our work suggests that this potential is enhanced when curli fibers are extracted from biofilms grown in environments with limited nutrients, as indicated by their higher polarity (**Figure 4a**). More generally, the water uptake of biofilms is regulated by the composition and characteristics of their extracellular matrix.^34^ Indeed, biofilm matrix influences the osmotic pressure driving water flows leading to nutrient transport and subsequent bacterial growth, but also to biofilm expansion via matrix swelling, as showed for B. subtilis and V. cholera.^34,35^ The trend of higher curli fiber yield when nutrients are less abundant is also consistent with bacteria having less metabolic and proliferation activities in such conditions (**Figure 22d** and **2e**). The nutrient concentration and more generally the composition of the substrate influence the composition of the biofilm matrix and subsequently biofilm properties.^4,9,11^ In B. subtilis biofilms for example, the composition of the growth medium was shown to actively contribute to the production, the distribution and the characteristics of the amyloid fibers found in the extracellular matrix.^4^ *V. fischeri* biofilms also acquire different morphologies and resistance to mechanical disruption depending on the nutrients available to the bacteria.^9^ Interestingly, bacteria grown on agar with the highest nutrient content (12% w/v) produced smaller biofilms containing less curli fibers (**Figure 11** and **2** ). This observation is consistent with the general understanding of matrix secretion and subsequent biofilm formation as part of the multiple bacteria responses to starvation.^32,36^

Biofilms can be considered as hydrogels and their mechanical properties are expected to be determined by both their network of biopolymers and their water content.^5^ Here we observe the counterintuitive result that biofilm stiffness is higher in conditions where biofilm water content is larger (**Figure 11d** and **2a**). This can be explained by a denser biopolymer network as indicated by the larger yield of curli fibers obtained in these conditions (**Figure 2f** ). Beyond their amount, curli fibers are likely to contribute to the macroscopic mechanical properties of the biofilms through their molecular structure (**Figure S12** ). Indeed, curli fibers are known for the relatively high rigidity provided by the cross-β sheet arrangement adopted by the CsgA subunit.^15,17,37^ Variations in the conformation of the amyloid proteins are thus expected to induce variations in the rigidity of the fibers, which in turn are expected to influence biofilm stiffness. Consistently, analyzing the structure of the curli fibers extracted from biofilms obtained in the various conditions revealed a relatively higher packing and β-sheet content, when obtained from the stiffest biofilms, i.e. at 1.5 % w/v nutrient concentration (**Figure 2d, 3b** and **3d,** and **Figure S9** ). We interpret the ThioT ratio of each fiber as fiber packing due to a combined interpretation of the ATR-FTIR spectroscopy, CD spectroscopy and intrinsic fluorescence analysis (**Figure 3d, Figure S9** and **Figure 4b** ). Specifically, the curli fibers that showed low ThioT intensity presented a low β-sheet content with ATR-FTIR and CD spectroscopy as well as change in the Trp environment. Combined, these results suggests the ThioT signal could be describing the fiber packing.^19^ These structural variations could also explain the different fluorescence emission spectra in the presence of Nile Red, and thus the different amyloid fiber polarity measured as a function of nutrient availability during biofilm growth (**Figure 4a** and **S10a**).^38^ The same reasoning can apply to the Trp amino acid present in the CsgA subunits, which intrinsic fluorescence indicates the hydrophobicity of its close environment (**Figure 4b**, and **S10b** ).^23^ Testing the integrity of the different samples in the presence of urea also revealed that the chemical stability of curli increases with fiber packing, including an additional influence of their β-sheet content (**Figure 4d-f**). Considering the mode of action of urea, this result is consistent with the idea that the access of urea to the peptide is easier in loosely packed fibers, thus facilitating amyloid denaturation.^14,39^

The multiple experiments presented in this work converge to the major finding that nutrient availability in the surroundings of the bacteria affects not only the production, but also the structure of the curli amyloid fibers they produce (**Figure 6**). As this result may appear surprising in light of the existing literature on the bacterial amyloid curli, it is important to highlight that the structural investigations reported here were performed on amyloid curli fibrillated *in vivo*, i.e. in the native biofilm micro-environment.^13^ The present physico-chemical context is thus significantly different, which may have a substantial impact on the resulting fiber conformation. For example, bacterial cell membranes, as well as the presence of lipopolysaccharides (LPS) may promote fiber aggregation.^13^ The matrix molecular crowding, especially the presence of extracellular DNA, is also contributing to the fibrillation process of amyloid fibers.^13^ Other environmental conditions, such as pH were also shown to influence this process (although *in vitro*).^15^

Considering the protective function of biofilm matrix in challenging environments, one can question whether these variations in curli structure and their implications at the biofilm scale are the result of incidental physico-chemical interactions during the fibrillation process, or whether *E. coli* bacteria actively tune this process to provide amyloid curli fibers with specific properties (e.g. polarity, chemical stability, stiffness) in order to support the survival of the community (e.g. nutrient storage and transport, water retention, physico-chemical shield).^32^ An attempt to address this point with further experiments indicates that these two hypotheses are not necessarily exclusive (**Figure S13** ). Indeed, after denaturing^40^ the curli fibers and repolymerizing the CsgA units in different media, the overall structural conformations of the original fibers obtained from biofilms grown in different conditions was retained (**Figure S13g and h** ). Bacteria may thus produce CsgA monomers that are slightly different (probably in structure) depending on the matrix fiber structure and properties they need to cope with the given environment. Nevertheless, curli fibers reconstituted from CsgA monomers purified from biofilms grown in abundance of nutrient (LB 6%w/v) also showed structural variations depending on the re-fibrillation conditions (**Figure S13g and h**). Although the present approach allowed us to establish a proof of feasibility, another approach yielding more purified curli fibers is required to efficiently tackle the related questions about CsgA structure, adaptability, and curli biogenesis.

In general, the fundamental outcomes of this work can have great implications for the development of both anti-biofilm strategies and biofilm-based materials. Indeed, we found out that *E. coli* form overall weaker biofilms when exposed to environments that are either very poor or very rich in nutrients, mainly by altering the amount, packing and conformation of the amyloid curli fibers constituting their extracellular matrix (**Figure 5** ).^36^ This information could possibly motivate an overfeeding of bacteria coupled with antibiotic treatments to prevent the formation of strong biofilms and simultaneously target the vulnerable bacteria.^41^ In contrast, the relatively more robust curli fibers harvested from biofilms formed by *E. coli* exposed to intermediate nutrient levels may represent building blocks with high potential for making amyloid-based materials.^42^ The possibility of tuning properties of the bacterial amyloids *via* the conditions of their genesis may even expand the range of possibilities in the emerging field of engineered living materials.^43^

## Experimental Section

### Bacterial strain and growth

The biofilm-forming bacterial strain *E. coli* K-12 W3110 was used throughout this study. Salt-free agar plates (15mm diameter) were prepared with 1.8% w/v of bacteriological grade agar−agar (Roth, 2266), supplemented with 0.75%, 1.5%, 3.0%, 6.0% or 12.0% w/v tryptone (Roth, 8952) and 0.25%, 0.5%, 1.0%, 2.0% and 4.0% w/v yeast extract (Roth, 2363), respectively. After agar pouring, the plates were left to dry for 10 minutes with the lid open and 10 minutes with the lid partially open to avoid future condensation. Each agar plate was left to rest for 48 hours at room temperature before bacteria seeding. A suspension of bacteria was prepared from a single colony and grown overnight in Luria−Bertani (LB) medium at 37 °C with shaking at 250 rpm. Each plate was inoculated with arrays of 9 drops of 5μL of bacterial suspension (OD600 ∼ 0.5 after 10x dilution). After inoculation, the excess water evaporated from the drops and left bacteria-rich dry traces of comparable sizes from 4 to 8mm diameter, depending on the growth condition. Biofilms were grown for 5 days in total (∼120h) inside an incubator at 28 °C. Monitoring the relative humidity in the incubator showed that it remains around 30 %RH.

### Biofilm imaging

Three biofilms per condition were imaged with a stereomicroscope (AxioZoomV.16, Zeiss, Germany) using the tiling function of the acquisition software (Zen 2.6 Blue edition, Zeiss, Germany). To estimate the biofilm size, at least 12 independent biofilms were used to calculate their area using the Fiji software.^44^

### Biofilm dry mass and water uptake

The water content and water uptake of the biofilms were determined by scrapping 7 biofilms per condition from the respective agar substrates after 5 days of growth (∼120 h). Biofilms were placed in plastic weighing boats, and oven-dried at 60 °C for 3 h. Wet and dry masses (M_wet_, M_dry_) were determined before and after drying weighing the biofilm masses in an analytical balance, sensing up to 3 decimal positions.^27^ To determine the water uptake (W ), we added Millipure water in excess (5 mL) to the biofilms harvested from each condition, covered them with aluminum foil to avoid evaporation and left overnight. The water excess was removed and the biofilm samples were weighed again (M_rewet_). The biofilms water content in each growth condition was estimated with **Eq. (2)**

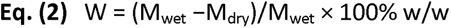

The percentage of water uptake of biofilms after rehydration *(%W _up_*) was determined with respect to the biofilm initial wet mass as described in **Eq. (3)**

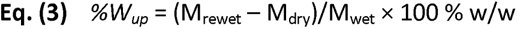

The water uptake per gram of dry biofilm *(W_up_*) was calculated with **Eq. (4)**

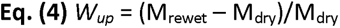

All procedures were carried out in four independent experiments.

### Microindentation on biofilms

Biofilms were grown for 5 days and stored at 4 °C, the Petri dishes were sealed with parafilm to prevent evaporation. A total of 4 biofilm samples were tested per condition. Microindentation measurements were carried out using a TI 950 Triboindenter instrument (Hysitron Inc.) to determine the load−displacement curves p−δ. The instrument was calibrated in air. Indentations were performed with a spherical diamond tip of radius R = 50 μm on a stage designed for the measurement of soft biological samples. The sample surface was approached up to 400 - 300 μm above the surface and retracted to the starting position while recording the measured force over the whole range. Loading rates ranged from 20 to 30 μm.s^-1^, which corresponds to loading and unloading times of 10 s. Between 8 and 10 measurements were performed in the central region of each biofilm, which were still attached to the respective agar substrates. Average and standard deviations were calculated over all measurements from a respective condition. The lateral distance between two measurement points was at least 200 μm in x and y directions. A Hertzian contact model was fitted to the loading part of the curve (indentation depth range δ = 0−10 μm) to obtain the reduced Young’s modulus (see **Figure S3** for more details).

### Biofilm total protein extraction and quantification

One biofilm per condition was scrapped and placed in an Epperndorf® tube with 200 µL of lysis buffer (1% SDS, 1% RIPA, 1% Tritonx100). The biofilms were disrupted mechanically using a XENOX MHX 68500 homogenizer for one minute, before being sonicated for 15 minutes in a bath sonicator. After sonication, biofilms were again disrupted for one minute and incubated at 26 °C for 20 minutes. The samples were then centrifuge at 17000 g for 10 minutes at 4 °C and the pellet was discarded. Aliquots of 50 µL were taken and mixed with an equal volume of an acetone/methanol 1:1 solution for protein precipitation. After an overnight incubation at - 20 °C, the samples were centrifuged at 18000 g for 30 minutes at 4 °C. The supernatant of each sample was discarded and pellets were resuspended in 15 µL of PBS 1X for protein quantification.

Protein quantification was done using a Bradford assay. Briefly, in a 96-well microplate, 5 µL of sample was incubated with 195 µL of Bradford reagent (Quick Start™ Bradford 1x Dye Reagent #5000205, BioRad). Absorbance was measured at 550 nm after a 5-10 minutes incubation protected from the light. BSA (heat shock fraction, protease free, fatty acid free, essentially globulin free, pH 7, ≥ 98%, (A7030) Sigma) was used in different concentrations as a calibration curve. The experiments were repeated with three independent biofilms.

### Bacterial growth curve

Salt-free Luria-Bertani liquid media were prepared at different nutrient concentrations for this experiment (0.75, 1.5, 3.0, 6.0 or 12.0 % w/v). Single colonies of *E. coli* W3110 K-12 bacteria were first grown on LB agar and were then inoculated in 10 mL of salt free LB media in 50 mL Falcon® tubes (one single colony per condition). The cultures were grown aerobically in a shaker (250 rpm) at 37 °C for 8 hours. Bacterial growth was monitored by measuring the OD_600nm_ of each culture every 30 minutes (see **Figure S4-5**). Three independent samples were tested for each condition.

The doubling time (T_d_) of each independent growth curve was calculated as **Eq. (5)**

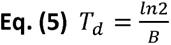

Where B is derived from the exponential fit of the linear part of the growth curve (see **Figure S4-5** for more details).

### Bacterial metabolic activity (MTT)

Salt-free Luria-Bertani liquid media were prepared containing different nutrient concentrations for this experiment (0.75, 1.5, 3.0, 6.0 or 12.0 % w/v). Single colonies of *E. coli* W3110 K-12 bacteria were first grown on LB agar and were then inoculated in 10 mL of salt free LB media in 50 mL Falcon® tubes (one single colony per condition). The cultures were grown aerobically in a shaker (250 rpm) at 37 °C for 8 hours. Aliquots of sample from the liquid cultures were taken at 60, 120, 240 and 360 minutes after the start point of the experiment to measure the metabolic activity of the bacteria. A protocol adapted from Oh et al.^45^ was used to perform an MTT (3-(4,5-dimethylthiazol-2-yl)-2,5-diphenyltetrazolium bromide) assay.

Briefly, 100 µL of sample were washed twice in PBS 1X by centrifugation at 1000 g for 5 minutes at 4 °C. The pellet was resuspended to a final volume of 100 µL PBS 1X. In a 96-well microplate, each sample was incubated with 10 µL of MTT 0.5 mg/mL in PBS for 15 minutes at 37 °C, protected from the light. After incubation, 100 µL of NaOH 1M was added to each sample in order to solubilize the formazan crystals. Absorbance values were recorded at 570 nm and PBS 1X was used as blank. The experiment was done with three independent bacteria liquid cultures per condition.

### Curli fiber purification

Fiber purification involved a similar process as reported in previous works.^14,46^ Briefly, a total of 27 biofilms (**∼** 1 g of biofilm material) were scraped from the surface of the substrates. Biofilms were blended five times on ice with an XENOX MHX 68500 homogenizer for 1 minute at 2-minute intervals. The bacteria were pelleted by centrifuging two times at low speed (5000 g at 4 °C for 10 min). A final concentration of NaCl 150 mM was added to the supernatant and the curli pelleted by centrifuging at 12.000 g at 4 °C for 10 minutes. The pellet was resuspended in 1 mL of solution containing 10 mM tris (pH 7.4) and 150 mM NaCl, and incubated on ice for 30 min before being centrifuged at 16.000 g at 4 °C for 10 minutes. This washing procedure was repeated thrice. The pellet was then resuspended in 1 mL of 10 mM tris solution (pH 7.4) and pelleted as described above (16.000 g at 4 °C for 10 minutes). The pellet was again suspended in 1 mL of 10 mM tris (pH 7.4) and centrifuged at 17.000 g at 4 °C for 10 minutes. This washing step was repeated twice. The pellet was then resuspended in 1 mL of SDS 1 % v/v solution and incubated for 30 min. The fibers were pelleted by centrifuging at 19.000 g at 4 °C for 15 min. The pellet was resuspended in 1 mL of Milli-Q water. This washing procedure was repeated thrice. The last resuspension was done in 0.1 mL of Milli-Q water supplemented with 0.02 % sodium azide. The fiber suspension was stored at 4 °C for later use. The protein concentration in monomeric units of the suspensions was determined by the absorbance from an aliquot incubated in 8 M urea at 25 °C for 2 h, a treatment leading to complete dissociation of the fibrils as verified by Thioflavin T measurements.

### Transmission electron microscopy (TEM)

2 μL drops of fiber suspension were adsorbed onto Formvar-coated carbon grids (200 mesh), washed with Milli-Q water, and stained with 1 % w/v uranyl acetate. The samples were imaged in a JEOL-ARM F200 transmission electron microscope (JEOL GmbH, Germany) equipped with two correctors for imaging and probing. For the observations, we applied an acceleration voltage of 200 kV.

### Attenuated total reflectance Fourier transform infrared spectroscopy (ATR-FTIR)

IR spectra were acquired with a spectrophotometer (Vertex 70v, Bruker Optik GmbH, Germany) equipped with a single reflection diamond reflectance accessory continuously purged with dry air to reduce water vapor distortions in the spectra. Fibers samples in Milli-Q water (∼10 μL) were spread on a diamond crystal surface, dried under N_2_ flow to obtain the protein spectra. A total of 64 accumulations were recorded at 25 °C using a nominal resolution of 4 cm^−1^.

Spectra were processed using Kinetic software developed by Dr. Erik Goormaghtigh at the Structure and Function of Membrane Biology Laboratory, Université Libre de Bruxelles, Brussels, Belgium. After subtraction of water vapor and side chain contributions, the spectra were baseline corrected and area normalized between 1700 and 1600 cm^−1^ (**Figure S7** ). For a better visualization of the overlapping components arising from the distinct structural elements, the spectra were deconvoluted using Lorentzian deconvolution factor with a full width at the half maximum (FWHM) of 20 cm^−1^ and a Gaussian apodization factor with a FWHM of 30 cm^−1^ to achieve a line narrowing factor K = 1.5.^21^ Second derivative was performed on the Fourier self-deconvoluted spectra for band assignment. The bands identified by both procedures were used as initial parameters for a least square iterative curve fitting of the original IR band (K = 1) in the amide I’ region, using mixed Gaussian/Lorentzian bands. Peak positions of each identified individual component were constrained within ±2 cm^−1^ of the initial value.

### Fluorescence spectroscopy

Corrected steady-state emission spectra were acquired with a FluoroMax®-4 spectrofluorometer (HORIBA). Spectra were recorded at 25 °C using a 3-mm path cuvette (Hellma® Analytics). Thioflavin T(ThioT) measurements were performed at final concentrations of 3 μM protein, 1 mM probe in Glycine buffer, pH 8.2, using λ_exc_ = 446 nm and spectral bandwidths of 10 nm. Nile red (NR) measurements were performed at a final concentration of 7 μM protein, 7 mM probe in water, using λ_exc_ = 560 nm and spectral bandwidths of 10 nm. Intrinsic fluorescence spectra (5 μM protein) were acquired using λ_exc_= 280 nm and 5/5 nm slit bandwidths. 5 µM solution of soluble Trp (N-Acetyl-L-tryptophan, NATA) was included as a reference of the emission spectrum of a fully exposed Trp.

### Chemical stability assay

To test the chemical stability of the protein fibers, 5 μM samples were prepared by incubation of increasing urea concentration (0 – 8 M) and left for 2 h at room temperature to ensure equilibrium. ThioT was then added in a final concentration of 1 mM and fluorescence emission spectra of the samples were acquired under excitation at λ_exc_ = 446 nm and spectral bandwidth of 5 nm. Emission of the ThioT in urea was done and no significant signal was detected. The IC50 value was estimated by fitting the data to a linear regression (Y = a * X + b) and then calculating IC50 = (0.5 - b)/a.

### Statistical analysis

For each experiment, 3 to 4 fiber solutions were used, where each solution came from different fiber purification batches. For each purification, 27 biofilms were cultured in each of the 5 growth conditions tested (i.e. on salt-free LB-agar plate containing 0.75 %, 1.5 %, 3.0 %, 6.0 % and 12.0 % w/v nutrient, respectively). For each batch of biofilm culture, the different samples of fibers obtained were treated simultaneously (or in consecutive days) to prevent variability due to unavoidable slight variations in the implementation of the protocols (e.g. temperature and humidity in the laboratory during agar preparation and/or biofilm seeding). For statistical analysis, a One-way ANOVA test was carried out. Mechanical properties data was analyzed using a Mann-Whitney U test. Unless otherwise stated in the caption, Dunn’s post-test for multiple comparisons were done with respect to the 1.5 % w/v nutrient salt-free LB-agar condition, considered as the standard seeding condition. Details of each test are described in the legend of the figures.

## Supporting information

Suuporting Information section

## Supporting Information

The following files are available free of charge:

- Table S1: Composition of salt-free LB agar substrates
- Figure S1: Characterization of the salt-free LB-agar substrates
- Figure S2: Biofilm morphology
- Figure S3: Microindentation loading curve
- Table S2: Composition of the *E. coli* W3110 biofilms grown on substrates of different nutrient concentration
- Figure S4: Biofilm mass
- Figure S5: Bacteria growth curve
- Figure S6: Bacteria in liquid media
- Figure S7: B Purified curli fiber TEM images
- Figure S8: ATR-FTIR spectra of the purified curli fibers
- Table S3: Secondary structure analysis of the purified curli fibers assessed by ATR-FTIR
- Figure S9: Fiber structure by circular dichroism (CD) spectroscopy
- Figure S10: Fiber polarity
- Figure S11: Curli fibers and the CsgA subunit
- Figure S12: Relationship structure/function between purified curli fibers and the biofilm mechanical properties
- Figure S13: Proof of concept: from purified curli fibers to CsgA to fibers again

## Data Availability Statement

The data that support the findings of this study are available from the corresponding authors upon reasonable request.

## AUTHOR INFORMATION

### Author Contributions

The manuscript was written through contributions of all authors. All authors have given approval to the final version of the manuscript. The authors declare no competing interests.

### CRediT author statement

**Conceptualization:** C.M.B and M.S. **Visualization:** C.M.B and M.S. **Methodology:** C.M.B and M.S. **Investigation:** M.S, and M.V.D. **Writing Original Draft:** M.S **– Review and Editing:** C.M.B and M.S. **Supervision:** C.M.B. **Project Administration:** C.M.B. **Resources:** C.M.B.

## ACKNOWLEDGMENT

M.S. acknowledges support from the Max Planck Queensland Centre on the Materials Science for Extracellular Matrices. The authors also thank Christine Pilz-Allen for her technical support in the laboratories and Heike Runge for her help in doing the transmission electronic microscopy experiments. The authors are grateful to Dr. Agustín Mangiarotti for the useful discussions throughout this study. The authors are also grateful to Regine Hengge (HU Berlin) for providing the *E. coli* strain W3110 and to Eric Goormaghtigh from the SFMB group at the Université Libre de Bruxelles for providing the Kinetics Software.

