## Supplementary material for "Nutrient availability influences *E. coli* biofilm properties and the structure of purified curli amyloid fibers": Suuporting Information section

### Supporting Information

| Salt-free agar characterization | Table S1 |
| --- | --- |
|  | Figure S1 |
| Biofilm morphology | Figure S2 |
| Microindentation loading curve | Figure S3 |
| Biofilm mass | Figure S4 |
| Bacteria growth curve | Figure S5 |
| Bacteria in liquid media | Figure S6 |
| Purified curli fiber TEM images | Figure S7 |
| ATR- FTIR spectra of the purified curli fibers | Table S2 |
|  | Figure S8 |
| Fiber structure by circular dichroism (CD) spectrometry | Figure S9 |
| Fiber polarity | Figure S10 |
| Curli fibers and the csgA subunit | Figure S11 |
| Structure/function relationship between purified curli fibers and biofilm mechanical properties | Figure S12 |
| Proof of concept: from purified curli fibers to CsgA to fibers again | Figures S13 |

1. Salt-free agar characterization
   a. **Experimental section**
      i. Agar substrate plate preparation

Salt-free agar plates (15 mm diameter) were prepared with 1.8 % w/v of bacteriological grade agar−agar (Roth, 2266), supplemented with different concentration of tryptone (Roth, 8952) and yeast extract (Roth, 2363) (**Table S1**). Each agar plate was left to rest for 48 hours before bacteria seeding to ensure the correct evaporation of possible water excess. After 5 days at 28 °C, the agar plates were ready for characterization.

Table S 1 Nominal nutrient contents respectively for the salt-free agar substrates in this work.

| Nominal agar concentration (w/v%) | 1.8 | | 1.8 | | 1.8 | | 1.8 | | 1.8 | |
| --- | --- | --- | --- | --- | --- | --- | --- | --- | --- | --- |
| Nutrient concentration (w/v%) | 0.75 | | 1.5 | | 3 | | 6 | | 12 | |
| Nutrient (g)/100 mL  *Tryptone : Yeast* | 0.5 | 0.25 | 1 | 0.5 | 2 | 1 | 4 | 2 | 8 | 4 |
| Effective water content (w/w%) | 95.03 | | 93.61 | | 90.84 | | 85.53 | | 75.75 | |

ii. Thermal Gravimetric Analysis (TGA)

TGA were carried out in TG 209 F1 Libra (Netzch-Gerätebau GmbH, Germany). Evaporation of water from gels was measured by heating ~10 mg of sample contained in alumina pans at a rate of 10.0 K.min^-1^ from 25 to 600 °C in nitrogen. Measurements were performed in triplicate.

iii. Microindentation of agar substrates

The microindentation experiments were carried out as described in Ziege et al.^1^ After the agar substrate preparation, 2-3 agar plates were used for microindentation. Seven measurements were performed on each sample. The distance between two measurement points was at least 250 μm in x and y directions and the depth of the indentation was between 10 and 30 µm. A TI 950 Triboindenter (Hysitron Inc.) was used to determine the load–displacement curves after calibration of the instrument in air. Loading rates ranged from 20 to 30 μm.s^-1^, which corresponds to loading and unloading times of 10 s. The loading portion of all curves were fitted with a Hertzian contact model over an indentation range of 0 to 10 µm to obtain the reduced Young’s modulus E_r_.

b. **Results**

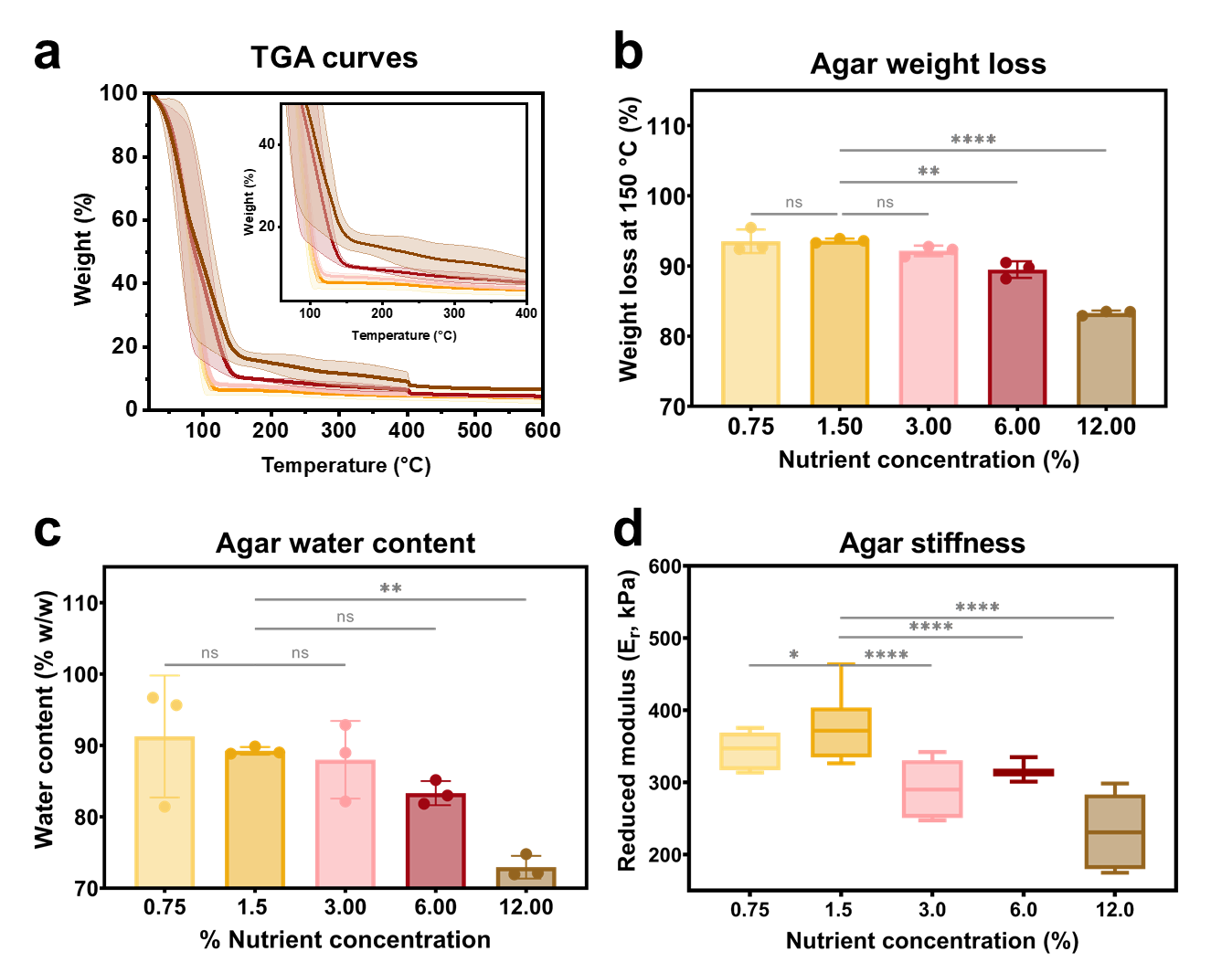

Figure S 1 Agar substrate properties. (a) Average Thermal Gravimetric Analysis (TGA) curves of each agar substrate between 90 and 600 °C. Inset is a zoom between 60 – 400 °C. N = 3. (b) Percentage of weight loss representing water loss at 150 °C for each agar substrate. (c) Water content of the agar substrates. N=3 .The statistical analysis was done with One-way ANOVA (p<0.0001, **** | p<0.001, *** | p<0.01, ** | p<0.05, * | ns = non-significant), where the 1.5 % w/v nutrient concentration condition was used as reference for the post-test multicomparisons. (d) Mechanical properties (stiffness) of the different agar substrates studied. N=14 microindentation curves of 2-3 agar substrates per condition. The number of individual measurements is n = 10−23 per condition. The statistical analysis was done with Man Whitney U test (p<0.00=1, **** |p<0.001, *** | p<0.01, ** | p<0.05, * | ns = non-significant), where the 1.5 % nutrient concentration condition was used as reference for the post-test multicomparisons.

We estimated the substrates (agar) free water content in every condition by carrying out thermal gravimetric analysis (TGA) (**Figure S1a**). We tested the behavior of the substrates from 25 to 600 °C. The results show that the higher the nutrient concentration in the agar substrate, the lower content of free water is in the substrate (**Figure S1a**). The loss of weakly linked water (Van der Waal forces) is indicated by a sharp loss of mass between 100 and 150 °C depending of the substrate. For substrates with low nutrient concentration, a lower temperature is needed to release weakly linked water compared to high nutrient concentration substrates. The total weight loss at 150 °C is correspond to the mass free water released (**Figure S1b**).^2^ Substrates with low nutrient concentration (0.75, 1.5 and 3.0 % w/v) lose ~90 % of their weight, while substrates with high nutrient concentration (6.0 and 12.0 % w/v) lose 83 and 73 % of their weight respectively. We also analyzed the water content of the salt-free LB-agar substrates by dehydration (**Figure S1c**). Low nutrient concentration substrates (0.75 and 1.5 % w/v) have 91 and 89 w/w % of water, respectively. Higher nutrient concentration substrates (3.0, 6.0 and 12.0 % w/v) have 87, 83 and 73 % w/w of water.

Analyzing the mechanical properties of the agar substrates revealed that the higher their nutrient content, the lower their stiffness (**Figure S1d-e**).

2. Biofilm morphology

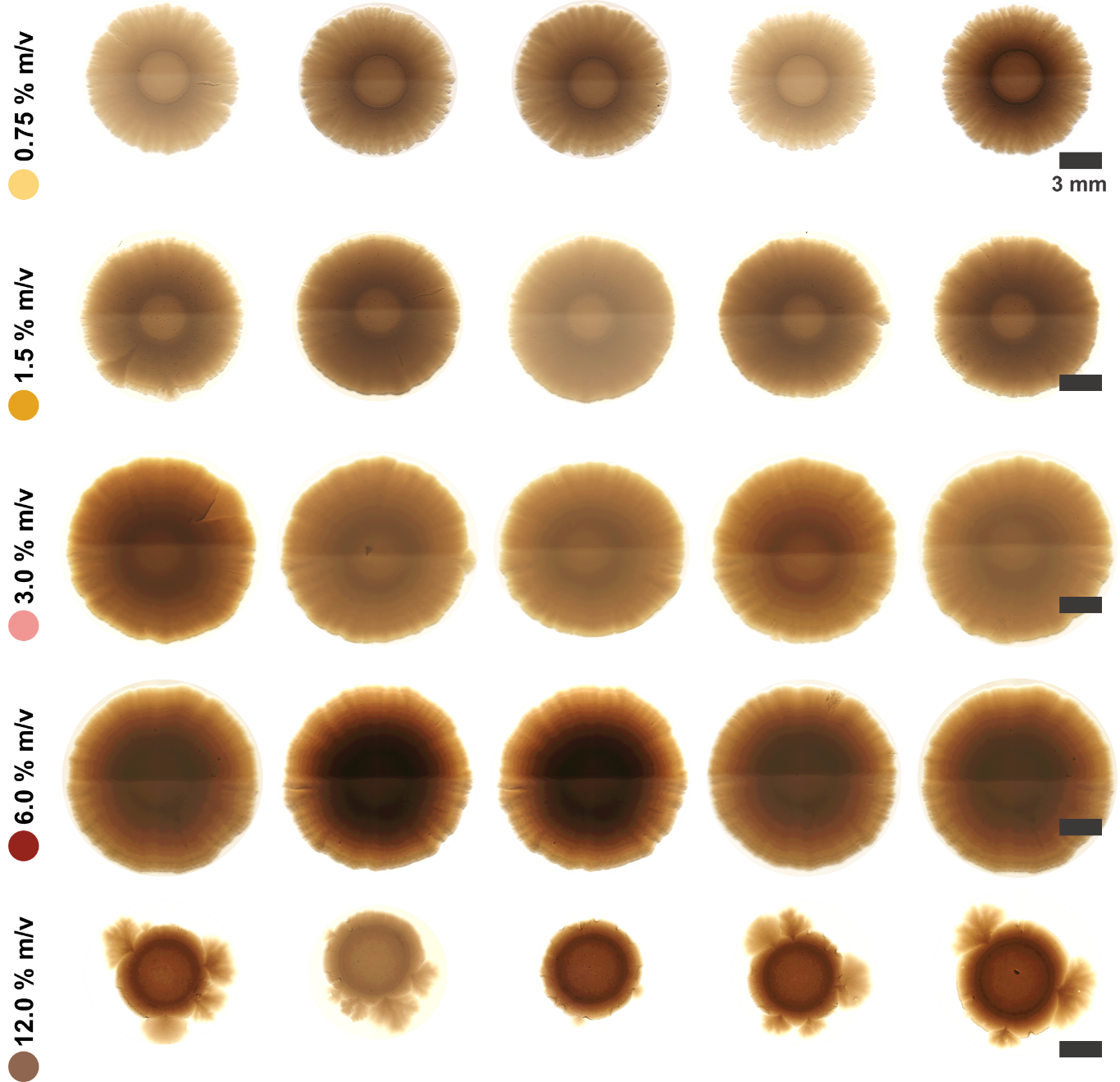

Figure S 2 Images from biofilms grown at different substrate nutrient concentrations. The images are representative of the different morphologies. Scale bar = 3 mm.

3. Biofilm mass

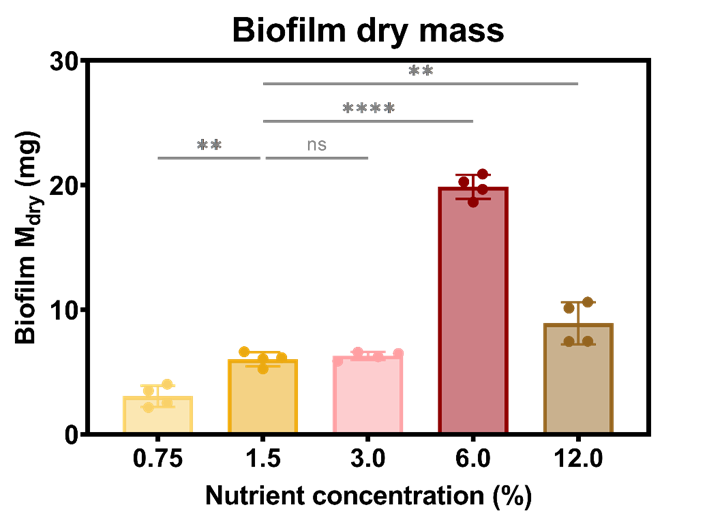

Figure S 3 Biofilm dry mass. Values correspond to the mass of a single biofilm calculated from four independent experiments. The statistical analysis was done with One-way ANOVA (p<0.0001, **** | p<0.001, *** | p<0.01, ** | p<0.05, * | ns = non-significant), where the 1.5 % w/v nutrient concentration condition was used as reference for the post-test multicomparisons.

4. Microindentation loading curves

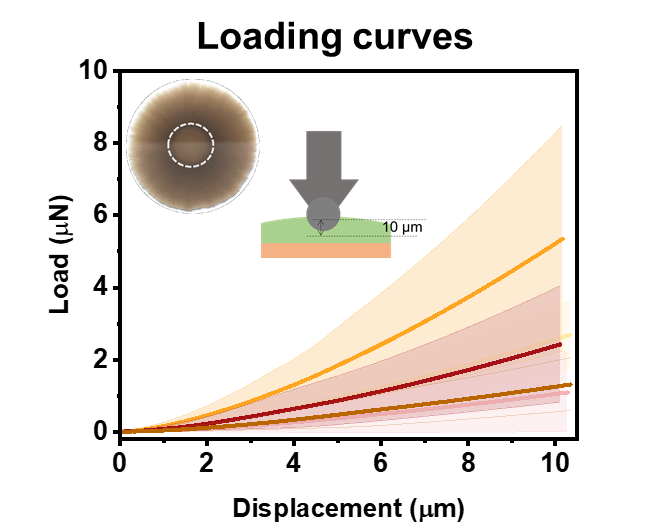

Figure S 4 Loading curves obtained from nanoindentation experiments on *E. coli* W3110 biofilms with different nutrient concentrations. Representative load−displacement curves when indenting the biofilm surface (loading curve). N= 40. Color code: 0.75 % w/v nutrient concentration (yellow), 1.50 % w/v nutrient concentration (orange), 3.00 % w/v nutrient concentration (pink), 6.00 % w/v nutrient concentration (red) and 12.0 % w/v nutrient concentration (brown). In the inset, a scheme depicts the measurement: a spherical tip (50 μm) was indented in ~10 μm of the biofilm to avoid contribution of the underlying salt-free LB agar. For more details see the experimental section.

5. Bacteria growth curve

Growth curves allow studying the kinetics of bacterial growth in specific media. Typically, these curves present the four phases of life for bacterial culture: lag, exponential, stationary, and death (**Figure S5**).^3^ In this work, the focus was set on the exponential phase, which is the phase where the bacteria have the most active metabolism.^4^

We can estimate the bacteria doubling time using the data from the exponential phase of the bacterial growth curve. The doubling time is the fastest rate at which the bacteria can replicate under the given conditions, and can be used to compare the effect of different growth media on the bacteria.^3^ The T_d_ is calculated by dividing the natural logarithm of 2 (ln2) by the exponent of growth (B) taken from the equation of the fitting curve: Y=Ae^Bx^.

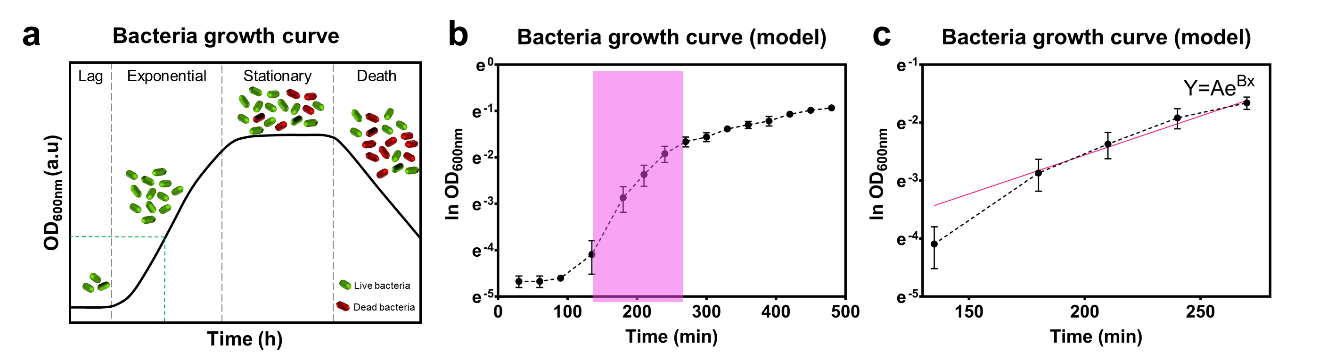

Figure S 5 Bacterial growth curve. (a) Bacteria grown in batch culture progress through four phases of growth: lag, exponential, stationary, and death. (b) Plotted growth curve ln(2) OD_600nm_ vs Time (min). Highlighted in pink is the linear section of the exponential phase. (c) Selected linear section of the exponential phase in the bacterial growth. A pink line represents the exponential regression fitting.

6. Bacteria in liquid media

In order to study whether the differences of protein production are due to the growth and metabolism of the bacteria, bacterial growth was monitored for 8 hours in liquid salt-free LB media and their doubling time (T_d_) calculated (**Figure S5**). According to the bacterial growth curves (**Figure S6a**), the duplication rates of bacteria were similar in all the conditions tested (**Figure S6b**). Interestingly, differences in the optical density were observed at the endpoint of the experiment, which directly correlates to the number of bacteria in the suspension (**Figure S6c**). The results suggest that the higher the nutrient concentration of the media, the higher the bacteria proliferation. Because the endpoint of our experiment was still at an early stationary phase (**Figure S5a, Figure S6a**), the optical density values acquired can be related to a majority of live bacteria.

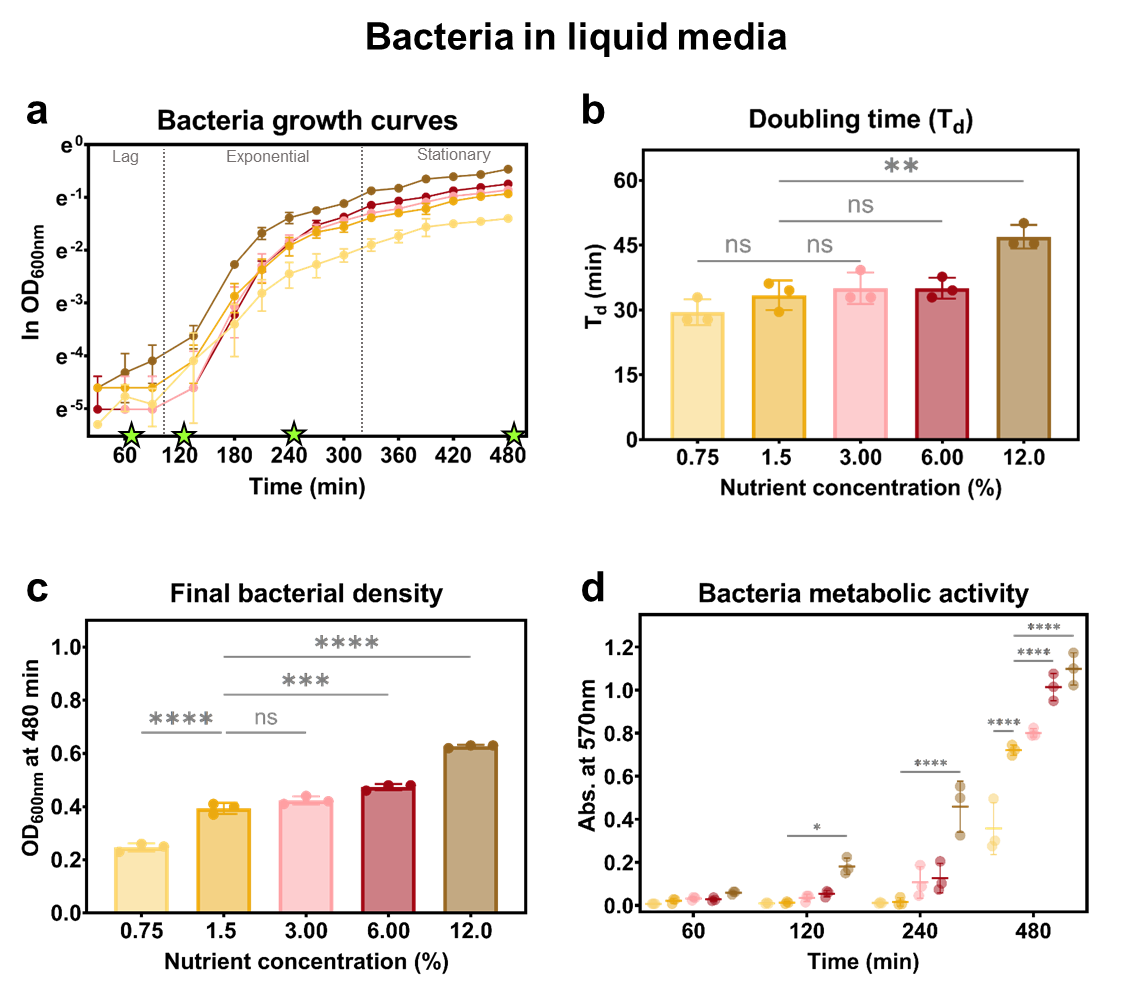

Figure S 6 Bacteria in liquid media. (a) 8-hour bacterial growth curves for each condition tested. (b) Bacteria doubling time (T_d_) calculated from each respective curve in (a). (c) Final optical density values acquired from each condition at the endpoint of the experiment. The optical density refers to the number of bacteria produced in each solution. (e) Bacteria metabolic activity in liquid media measured using MTT at the times highlighted by green stars in panel a. All data presented here come from N = 3 independent biofilm cultures for each condition tested. The statistical analysis was done with One-way ANOVA (p<0.0001, **** | p<0.001, *** | p<0.01, ** | p<0.05, * | ns = non-significant), where the 1.5 % w/v nutrient concentration condition was used as reference.

7. TEM of the purified fibers

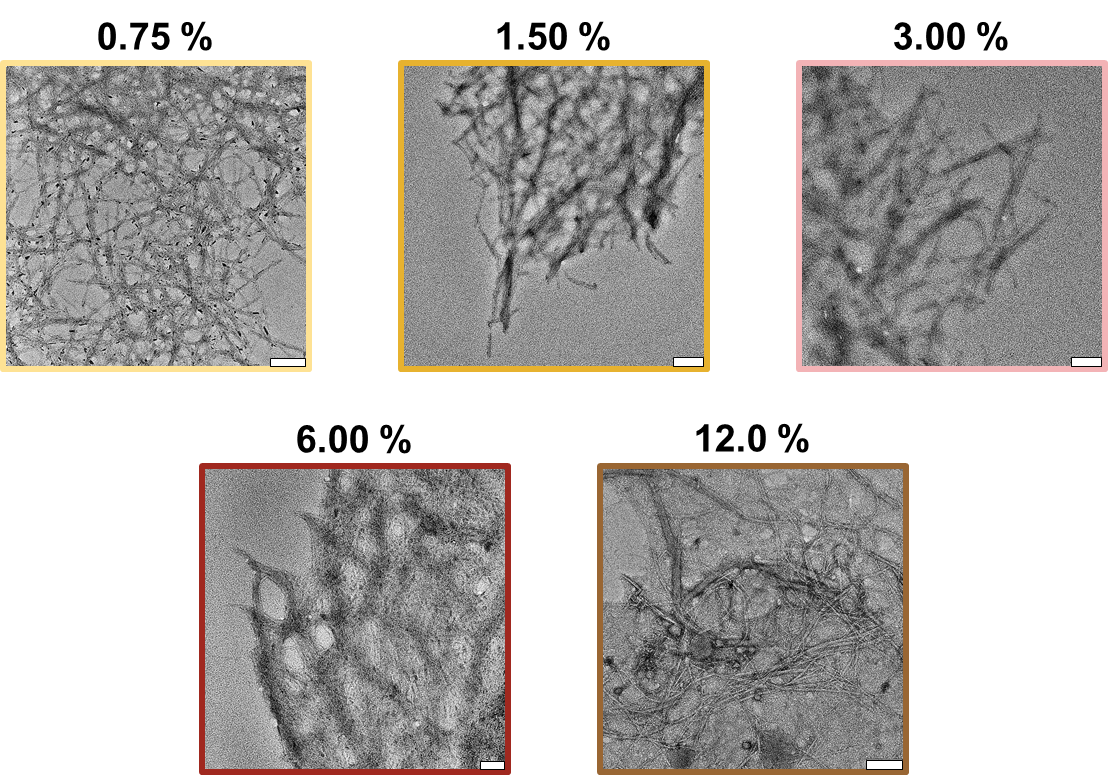

Figure S 7 Transmission electronic microscopy images of the purified fibers. Scale bar = 100 nm. Data come from 4-5 independent biofilm cultures for each condition tested.

8. FTIR-spectra of purified curli fibers

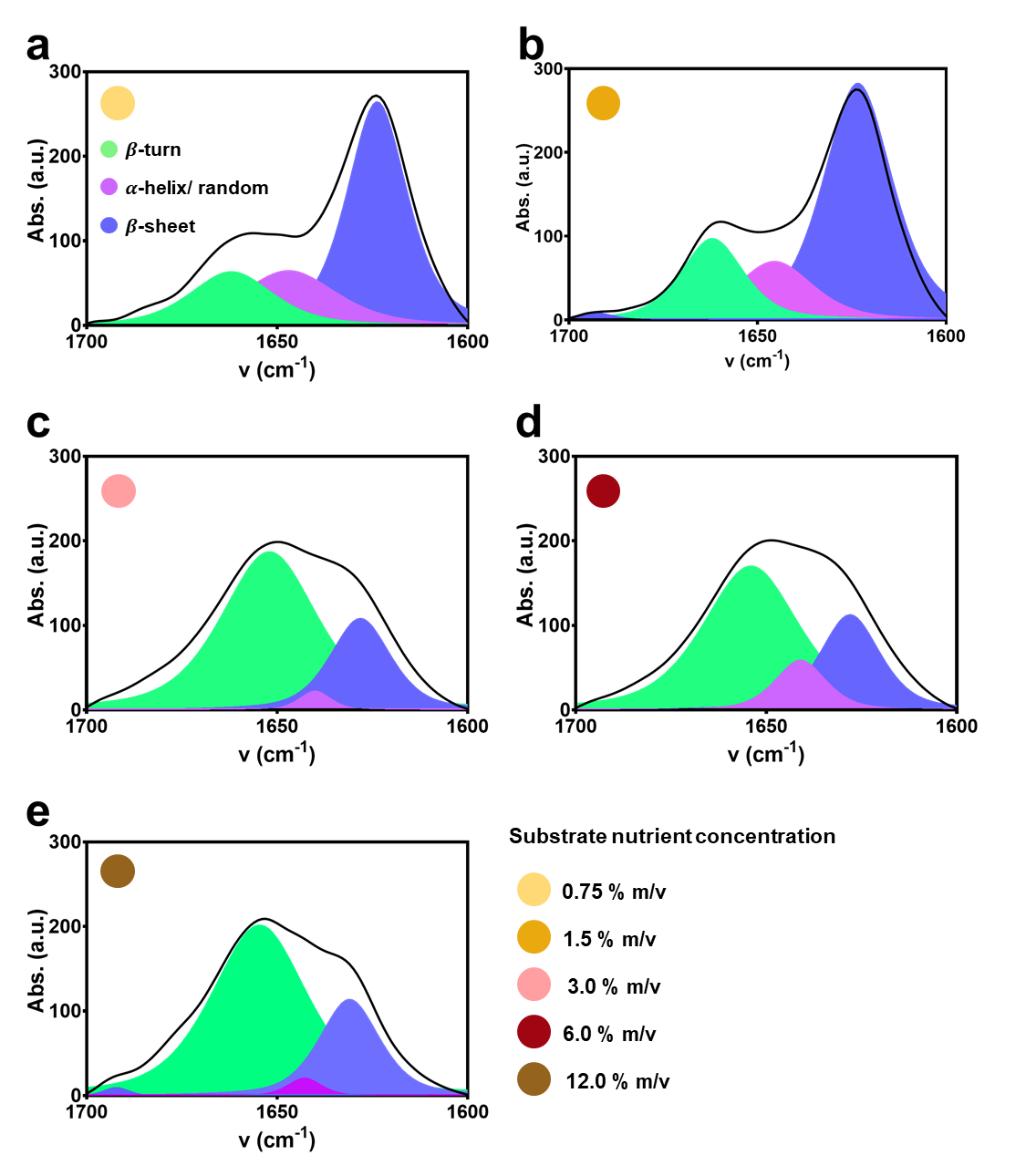

Figure S 8 Amide I’ spectra of curli fibers purified from *E. coli* biofilms. Area-normalized FTIR absorbance spectra in the Amide I’ region of the different samples. The spectra were curve-fitted using the number of spectral components identified by second derivative on the Fourier self-deconvoluted spectra. N = 3.

Table S 3 Secondary structure analysis of the purified curli fibers assessed by ATR-FTIR. The values correspond to Figure 3d. N = 3.

| Substrate nutrient concentration (% w/v) | Secondary structure content (%) | | |
| --- | --- | --- | --- |
|  | β - sheet | α - helix/ random | Turns |
| 0.75 | 58.03 ± 0.41 | 22.44 ± 0.12 | 19.82 ± 0.10 |
| 1.50 | 57.33 ± 1.04 | 29.63 ± 3.84 | 16.08 ± 1.63 |
| 3.00 | 30.08 ± 5.98 | 4.87 ± 3.00 | 65.01 ± 5.74 |
| 6.00 | 25.23 ± 3.66 | 14.84 ± 10.41 | 59.82 ± 6.92 |
| 12.00 | 26.72 ± 1.88 | 3.41 ± 0.78 | 69.86 ± 2.61 |

9. Fiber structure by circular dichroism (CD) spectrometry
   a. **Experimental section**

Spectra of 5µM CsgA monomer concentration of fiber solution in Milli-Q water were recorded with a Chirascan CD spectrometer (Applied Photophysics, Leatherhead, Surrey, UK). A quartz cuvette with 1 mm path length (Hellma, Müllheim, Germany) was used. Spectra are acquired between 190 nm and 250 nm wavelengths, with 1 nm step size, 1 nm band-width and 0.7 s integration time per point. Milli-Q water was used to define the measurement background, which was automatically subtracted during acquisition. The experiments were repeated thrice for each condition. Each measurement was an average of three scans.

b. **Results**

The purified curli fibers showed differences in their β-sheet structure as described in their CD spectra (**Figure S9**). All fibers tested in this experiment were normalized by CsgA monomer concentration, meaning there is the same protein concentration in each sample tested. Because of this, lower signal intensity means lower β-sheet content. Hence, fibers from biofilms grown on agar substrates containing lower nutrient concentration (0.75 and 1.5 % w/v), present high β-sheet content than those from biofilms grown on agar substrates containing higher nutrient concentration (3.0 % w/v or higher). These results are complementary to those of acquired with the ATR-FTIR experiments.

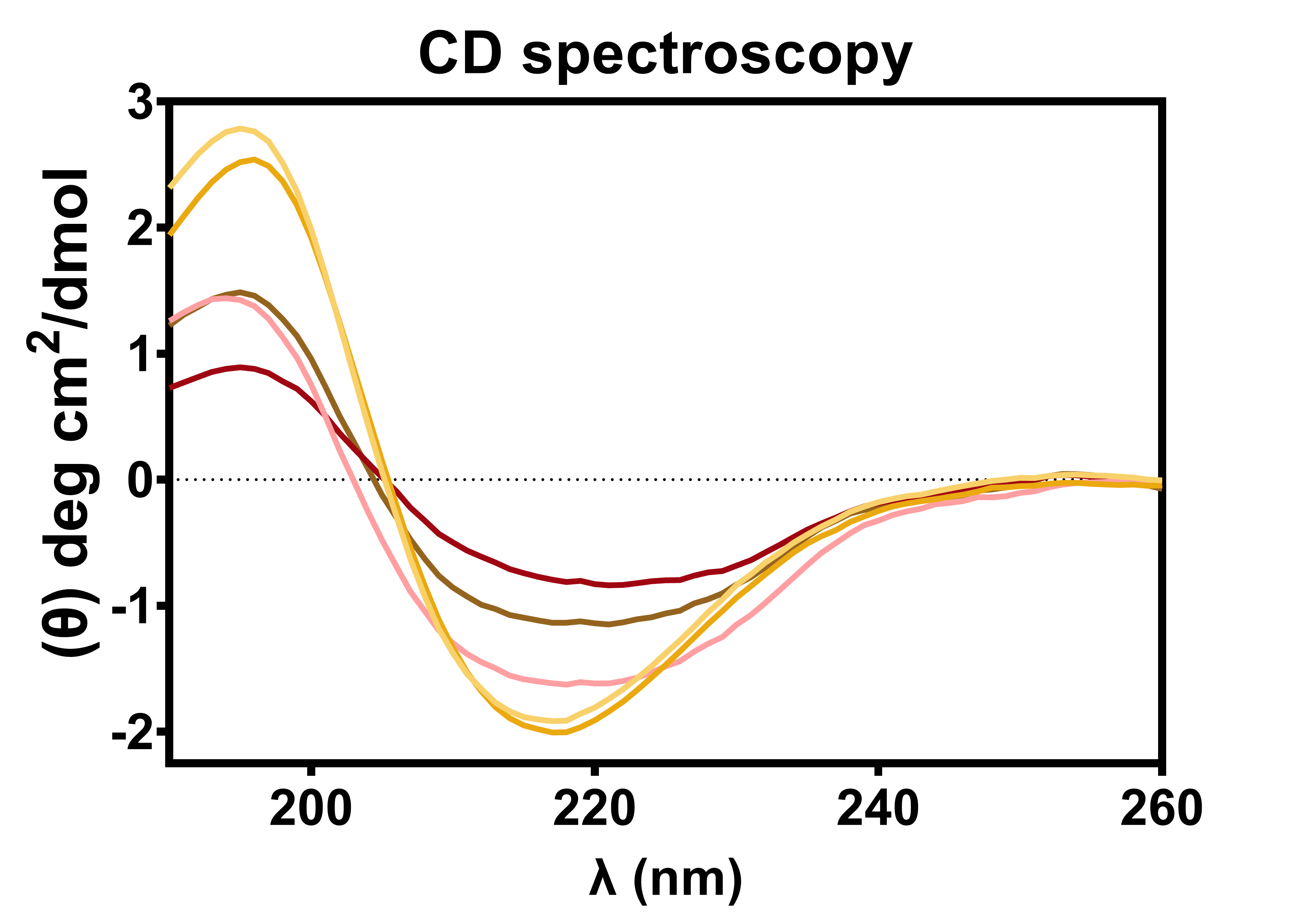

Figure S 9 CD spectrometry of the purified curli fibers. All fibers were normalized by CsgA monomer concentration. Data come from N = 3 independent biofilm cultures for each condition tested.

10. Fiber polarity

Fiber polarity is defined as the ratio between the position of the emission maximum of the nile red (NR) in buffer and the position of the emission maximum of the NR bound to each fiber (**Eq. S1**):

**Eq. (S1)** $Fiber polarity=\frac{Position of emission maximum {NR}_{fiber} (nm)}{Position of emission maximum {NR}_{buffer} (nm)}$

A polarity value of 1.00 indicates highly polar fibers.

Trp hydrophobicity is defined as the ratio between the position of the emission maximum of the Trp population in the fibers and the position of the emission maximum of soluble Trp (NATA) in buffer (as reference for the position of the emission maximum when we have the highest exposure to the solvent) (**Eq. S2**):

**Eq. (S2)** $Trp hydrophobicity=\frac{Position of emission maximum {Trp}_{fiber} (nm)}{Position of emission maximum {NATA}_{buffer} (nm)}$

A hydrophobicity value of 1.00 indicates more exposure of the Trp to the solvent.

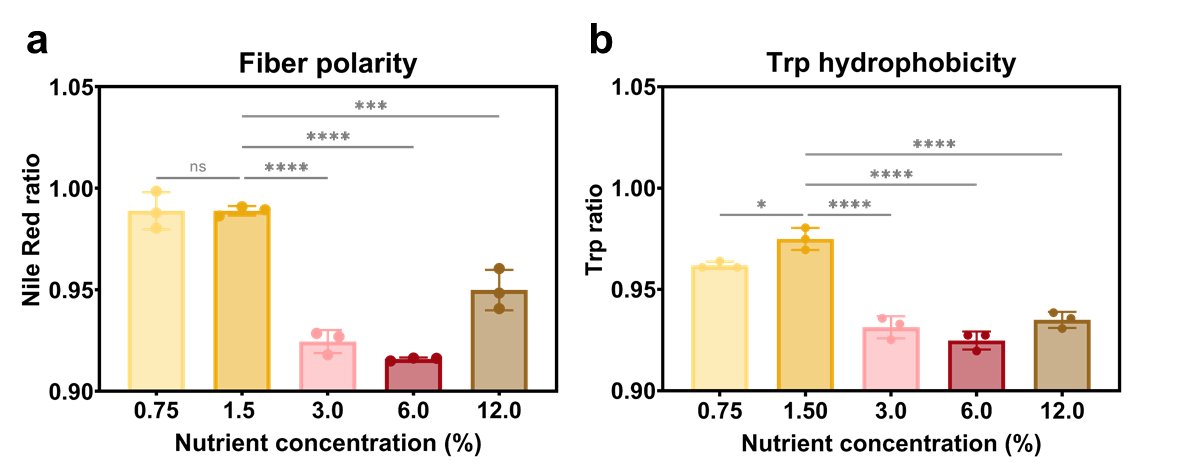

Figure S 10 Fiber polarity. (a) Ratio between the position emission maximum of each condition, taking NR in buffer as a reference. (b) Ratio between the position emission maximum of each condition, taking NATA in buffer as a reference. Data come from N = 3 independent biofilm cultures for each condition tested. The statistical analysis was done with One-way ANOVA (p<0.0001, **** | p<0.001, *** | p<0.01, ** | p<0.05, * | ns = non-significant), where the 1.5 % w/v nutrient concentration condition was used as reference for the post-test multicomparisons.

11. Curli fiber and CsgA subunit

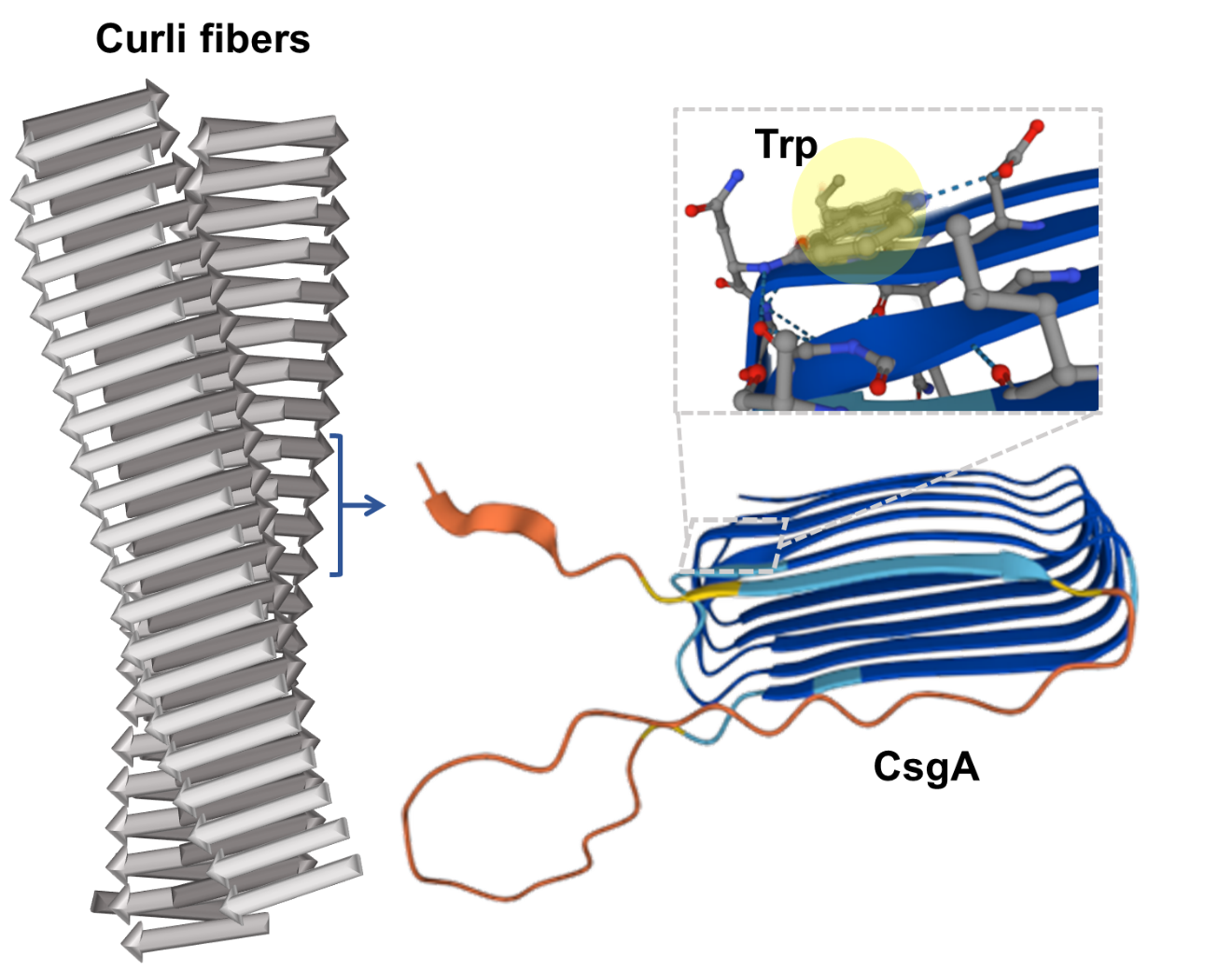

Figure S 11 Scheme of the composition of curli fibers and the position of the Trp in the CsgA protein.

12. Structure/function relationship between purified curli fibers and biofilm mechanical properties

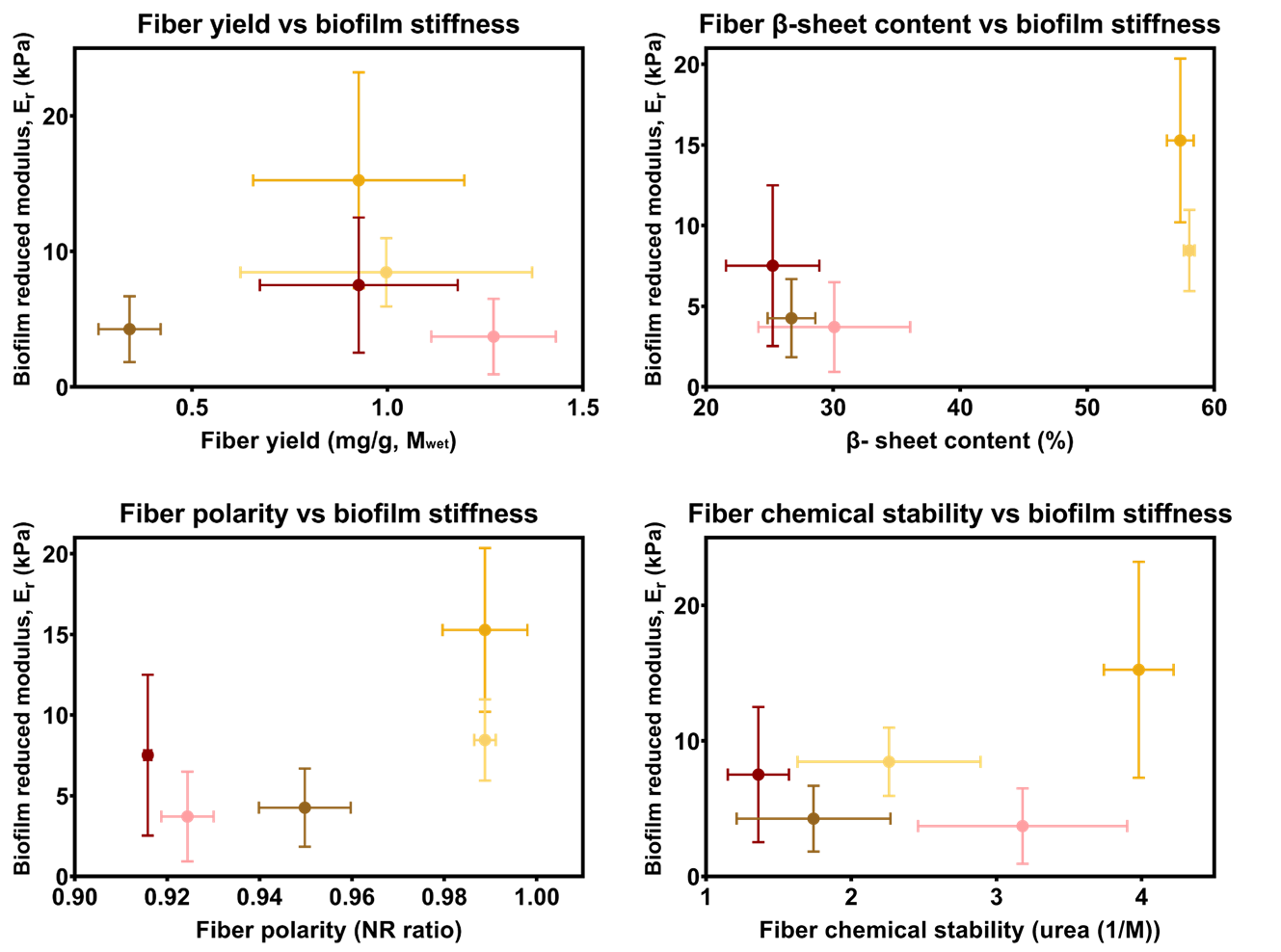

**Figure S 13 Biofilm mechanical properties as a function of different properties of the purified curli fibers.**

13. Proof of concept: from purified curli fibers to CsgA to fibers again
    a. **Experimental section**
       i. Purified curli fiber denaturation and CsgA polymerization

Samples of purified curli fibers were precipitated by centrifugation for 25 minutes at 18000g at 4 ºC. In order to denature the purified curli fibers, 20 µL of formic acid (FA) at 98% was added to each sample and left to evaporate after 15 minutes.^5^ After denaturation, 60 µL of milliQ water was added to the CsgA monomers and the samples were immediately prepared for polymerization in 96-well microplates (**Figure S13**).

Each well contained a final volume of 50 µL made of 10 µL of CsgA monomer solution, 5 µL of ThioT 100 µM and 35 µL of solvent. The solvent was either Tris-HCl 10 mM NaCl 50 mM, LB medium containing 1.5% w/v, or LB medium containing 6.0% w/v. The fibrillation of CsgA was monitored by placing the microplate in a microplate reader (BioTek Cytation5, Agilent), where the ThioT emission was acquired every 10 minutes for 10 hours. The temperature was set at 25 ºC and the plate was shaken prior to every measurement. The fibrillation process was performed and studied 4 times for each condition.

The CsgA monomers from the denatured samples were identified with 15% acrylamide SDS-PAGE electrophoresis gel, and CD spectroscopy. The polymerized CsgA fibers were identified using ATR-FTIR spectroscopy, transmission electronic (TE) microscopy, and ThioT fluorescence emission. These measurements were done twice.

b. **Results**

We purified the curli fibers from biofilms grown in two different conditions of the biofilm: LB agar substrates containing 1.5% w/v and 6.0% w/v nutrients. We then reduced these fibers to their monomer state (CsgA), and fibrillated them again in various buffers (**Figure S13**). To favor clarity, the curli fibers purified from biofilms grown on LB agar substrates containing 1.5% w/v nutrients are “Fiber A” and the curli fibers purified from biofilms grown on LB agar substrates containing 6.0% w/v nutrients are “Fiber B”.

CsgA monomers or polymers were identified by a band of the SDS-PAGE around 15 kDa and two bands around 30 kDa^6^, as well as by CD spectroscopy with a peak close to 200 nm^6^ (**Figure S13 b** and **c**). The increase of ThioT emission indicates that the previously denatured fibers can repolymerize in each of the media tested (**Figure S13d**). Fibers grown in LB 1.5% w/v show a slower polymerization process for the monomers of fiber A and fiber B compared to the other conditions tested. Fibers grown in buffer or LB 6.0% w/v have similar polymerization curves.

At the end of the polymerization experiment, each sample was centrifuged to separate the resulting fibers from the unpolymerized CsgA (supernatant) and restained with ThioT for fiber identification (**Figure S13e**). Because of the low yield in each sample, the concentration of fibers measured by ThioT emission was not normalized to any concentration. In all cases, the ThioT emission spectra observed displayed the shape expected in the presence of amyloid fibers in the sample. Moreover, electron microscopy (TEM) confirmed the existence of fibers (**Figure S13f**).

The ATR-FTIR spectra of the newly polymerized fibers in each medium showed spectra with peaks in similar positions to those of their respective original fibers (before denaturation with FA) (**Figure S13g**). Upon detailed analysis of these spectra, band assignment and secondary structure fitting suggest the structure of the newly polymerized fibers in each media follow the same trend as the structure of their respective original fibers (Fiber A and Fiber B, respectively), amidst the statistical differences observed (**Figure S13h**). These exploratory results suggest that depending on the environmental cues bacteria are under, they might express the CsgA monomer with a different secondary structure, thus defining different structural conformations of the mature curli fibers.

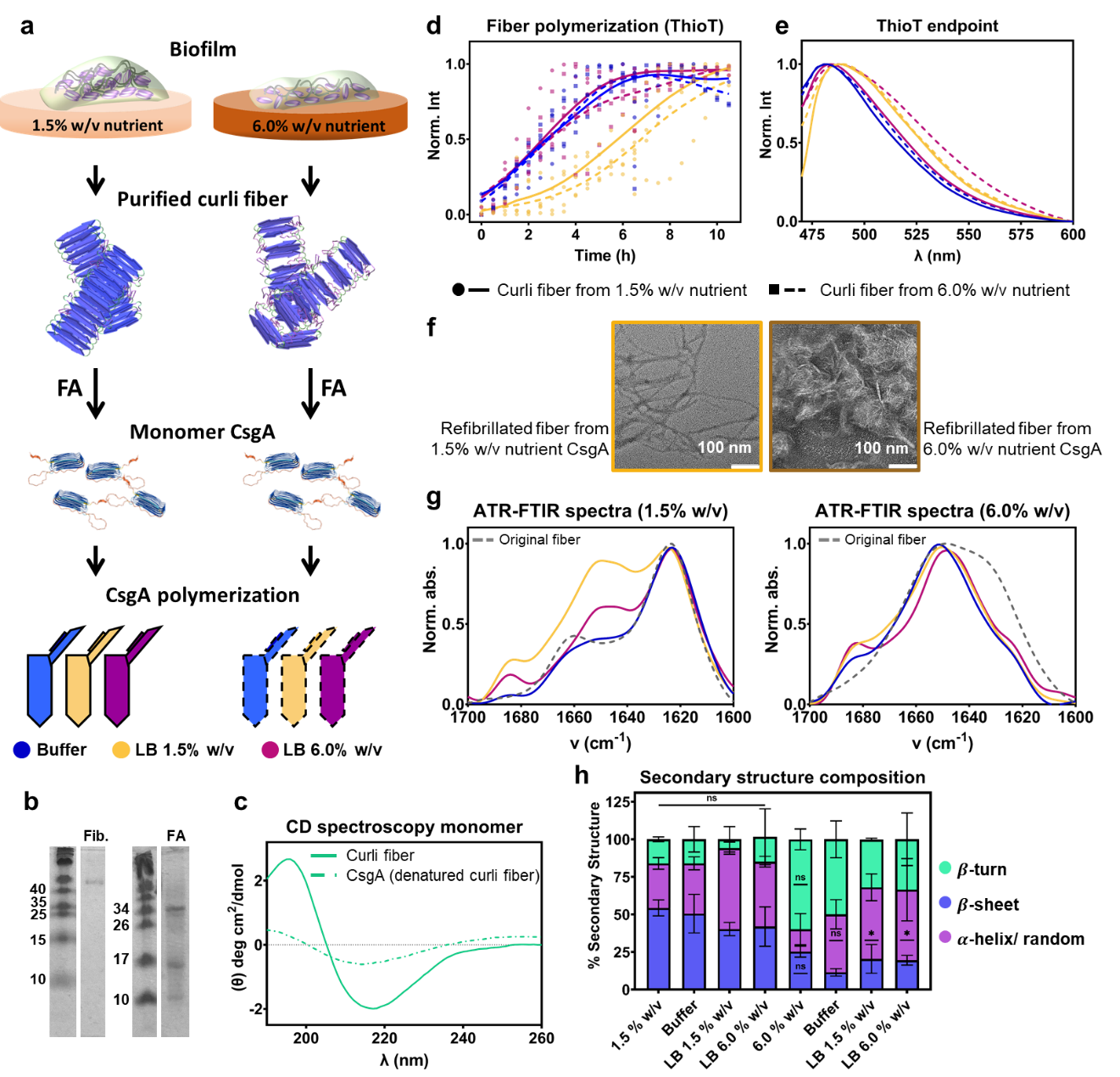

**Figure S 14 Curli fiber denaturation and characterization of refibrillated fibers.** (a) Schematic work-flow. (b) Coomassie stain SDS-PAGE purified curli fibers (Fib.) and of samples after fiber denaturation with formic acid (FA). Molecular size markers (in kDa) are indicated in the corresponding ladder. (c) CD spectra of the purified curli fibers and of the CsgA monomer after denaturation. (d) CsgA polymerization followed by ThioT emission. (e) ThioT fluorescence emission of the dye bound to the fibers after CsgA polymerization in (d). (f) TEM images of the polymerized fibers. Scale bar = 100 nm. (g) Amide I’ spectra of polymerized fibers. Area-normalized FTIR absorbance spectra in the Amide I’ region of the different samples. The spectra were curve-fitted using the number of spectral components identified by second derivative on the Fourier self-deconvoluted spectra. (h) Distribution of the three types of secondary structure in the polymerized fibers. The data was obtained from the Amide I’ region of each spectra. The statistical analysis was done with One-way ANOVA (p<0.0001, **** | p<0.001, *** | p<0.01, ** | p<0.05, * | ns = non-significant), where the 1.5 % w/v nutrient concentration condition was used as reference for the post-test multicomparisons. All data came from two independent denaturation experiments, except fiber polymerization that was done four times for each condition (duplicate of each denaturation experiment).
